# Sequential Design of Experiments enables accessible and efficient multidimensional mapping of single- and multi-protein condensates

**DOI:** 10.64898/2026.09.29.755056

**Authors:** Matej Siketanc, Ferdinand Weidner, Casimir Nüsperling, Maria Hondele

## Abstract

Biomolecular condensation emerges from networks of multivalent interactions that respond collectively to molecular composition and environmental conditions. Here, we establish a sequential Design of Experiments (DoE) framework to systematically and efficiently characterise multidimensional condensation landscapes. Applied to three RNA-binding condensate proteins – Dhh1, Ded1 and PGL-3 – the workflow revealed unexpected protein-specific interactions between biophysical and biochemical variables, including RNA, pH and ionic strength, and uncovered context-dependent switch-like behaviours. Extending the framework to protein mixtures revealed how proteins affected each other’s condensation and whether they formed condensate-within-condensate structures under different experimental conditions. Finally, an unexpected response identified by DoE prompted mechanistic analysis of Dhh1, revealing an ATP-independent RNA-binding surface that promotes efficient condensation in vitro and contributes to P-body formation in cells. Sequential DoE therefore provides an experimentally accessible strategy for uncovering context-dependent condensate regulation and converting multidimensional response patterns into mechanistic hypotheses.

## INTRODUCTION

Biomolecular condensates organise cellular biochemistry by concentrating selected proteins and nucleic acids into membraneless compartments. These assemblies contribute to diverse processes, including RNA metabolism, stress adaptation, signalling and developmental regulation^1–3^. Biomolecular condensate formation is driven by multivalent, transient interactions among proteins and nucleic acids^4–6^. Because these interaction networks are highly sensitive to molecular composition and the surrounding chemical environment, condensate formation is rarely controlled by a single variable. Variables such as protein and RNA concentration^7–9^, RNA sequence^10^, molecular crowding^11^, pH^7,12^, ionic strength^13^, temperature^14^ and nucleotide availability^15,16^, among many others^17^, can all influence whether condensates form, dissolve or change their internal organization. Importantly, these variables do not act independently: condensate behaviour emerges from their combined effects, such that the effect of one variable depends on the surrounding biochemical conditions. Condensate behaviour can therefore be highly context dependent^18–21^. Systematically characterizing this behaviour across combinations of influencing factors is essential to distinguish general regulatory principles from those specific to particular conditions.

This multivariable regulation presents an experimental challenge^9^. *In vitro* condensate studies commonly examine one factor at a time (OFAT), for example by titrating protein, RNA or salt concentration while keeping all other variables constant^13,22,23^. Such experiments are intuitive and well suited for testing defined hypotheses, but they only sample a narrow trajectory through a multidimensional experimental space. They may therefore miss conditions in which variables act together, overestimate effects that depend on a fixed background condition, or identify apparent local optima that disappear once additional variables are considered^24^. Understanding how condensates respond to their biochemical environment therefore requires approaches that systematically examine the combinatorial behaviour of several biochemical variables.

High-throughput phase-mapping platforms, including droplet microfluidics and related combinatorial approaches, provide powerful ways to sample multidimensional parameter spaces^25–31^. These methods can generate dense phase diagrams, but often require substantial amounts of protein, fluid-handling infrastructure and specialised analysis workflows. A complementary approach is therefore needed for experiments in which material, throughput or instrumentation is limited. Rather than maximizing the density of measurements across an entire parameter space, such an approach should prioritize informative combinations of conditions, distinguish the effects of individual variables from those that emerge in combination, capture nonlinear behaviour, and remain compatible with standard laboratory imaging assays.

Design of Experiments (DoE) provides a framework that addresses these challenges. Widely used in engineering, chemistry and process optimization, DoE applies statistically structured combinations of conditions to identify influential factors, quantify interactions and construct empirical models^32–36^. (Fig. 1a). An initial screening stage prioritizes the most influential factors, thereby reducing the number factors carried forward into more experimentally demanding designs. Subsequent full factorial designs quantify how these selected factors act individually and in combination. Finally, response-surface designs resolve nonlinear behaviour within the selected parameter space (Fig. 1b, c). This progression transforms multidimensional mapping into an experimental decision process, enabling the identification of informative response regions for independent validation and mechanistic investigation.

**Fig. 1.**
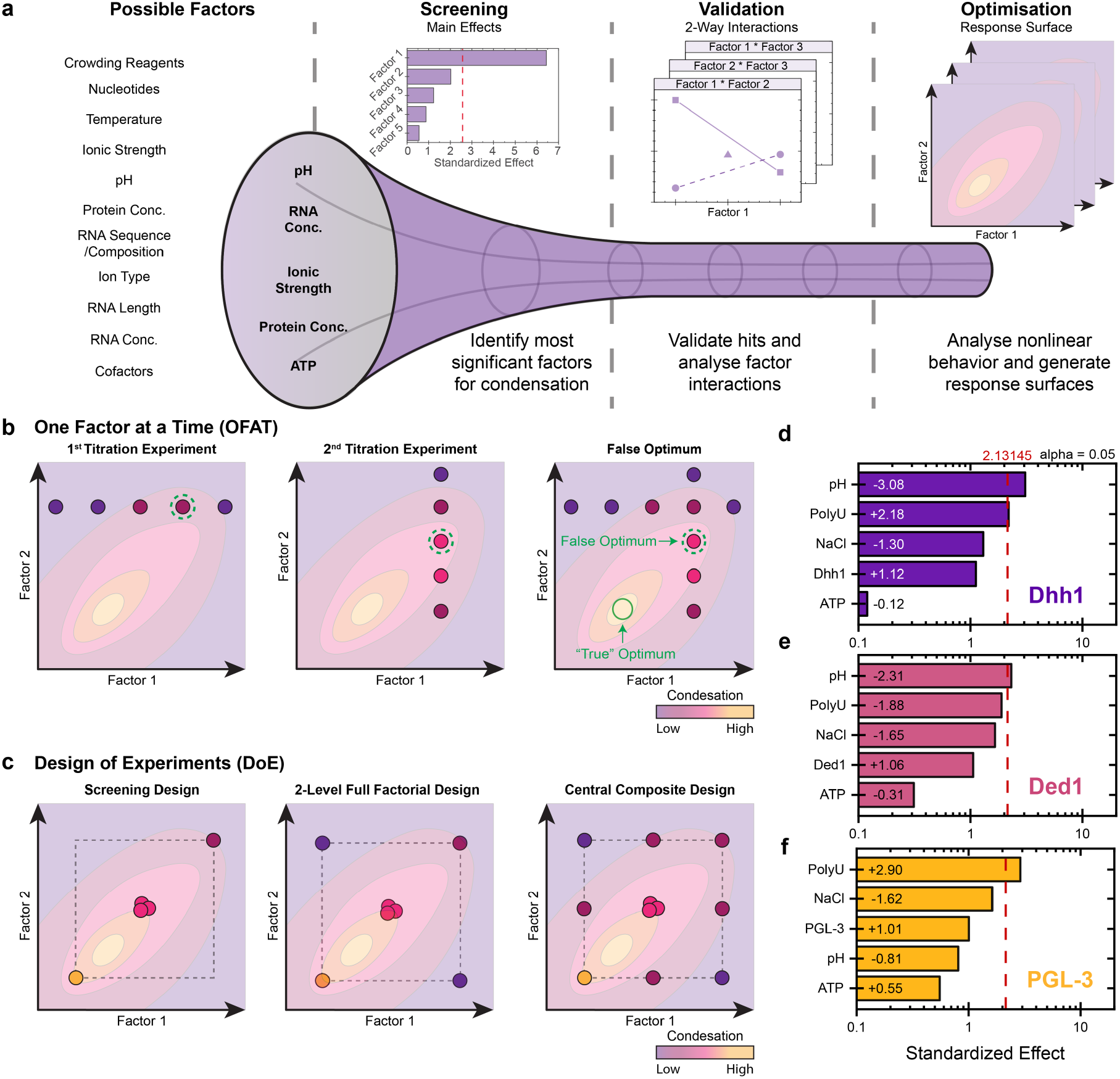
| Design-of-experiments workflow identifies protein-specific factors influencing condensate formation. **a** Schematic overview of the Design-of-Experiments (DoE) workflow used to characterise condensate behaviour. Five experimentally controlled factors (pH, RNA concentration, ionic strength, protein concentration and ATP concentration) were selected for initial screening, followed by higher-resolution designs to resolve interactions, nonlinear responses and multidimensional condensation landscapes. **b** Schematic comparison with a conventional one-factor-at-a-time (OFAT) strategy, which samples only restricted regions of the parameter space and may fail to identify response optima when factors interact. **c** Schematic representation of multivariable DoE approaches, including screening, full-factorial and response-surface designs, which progressively enable estimation of main effects, factor interactions and nonlinear responses. **d–f** Definitive screening design (DSD) analysis of Dhh1, Ded1 and PGL-3 condensation. ATP (0–4 mM), NaCl (100–200 mM), poly(U) (0–100 µg/mL), pH (6.5–7.5) and protein concentration (5–10 µM) were varied across an 11-condition definitive screening design. Bar length and the value displayed within each bar indicate the magnitude of the standardised effect. Positive values indicate that increasing the factor increased CV and therefore promoted condensation, whereas negative values indicate that increasing the factor decreased CV and therefore reduced condensation. The red dashed line marks the two-sided α = 0.05 significance threshold.

Here, we established a sequential Design of Experiments workflow for mapping multidimensional condensate behaviour using standard biochemical reconstitution and fluorescence microscopy. We applied this workflow to three model condensate proteins: the *Saccharomyces cerevisiae* DEAD-box ATPases Dhh1^16,37^ and Ded1^14,16,38^, and the *Caenorhabditis elegans* P-granule protein PGL-3^39,40^. Across all three systems, RNA, pH and ionic strength emerged as influential factors within the tested parameter space, but their effects were context dependent and integrated according to protein-specific regulatory logic. Extending the framework to mixed-protein systems showed that effects on bulk condensation and internal organization were distinct and protein-pair specific. Unexpectedly, the multidimensional maps revealed an unexpected regime in which Dhh1 formed RNA-containing condensates without added ATP. This observation prompted us to investigate a secondary RNA-binding surface and link it to Dhh1 condensation *in vitro* and P-body assembly in yeast cells. Thus, sequential multidimensional mapping can not only resolve context-dependent condensation behaviour but also uncover unexpected biological phenomena that motivate mechanistic investigation.

## RESULTS

### Screening identifies influential condensation factors

To establish an efficient workflow for systematically characterising condensate behaviour in vitro, we selected three RNA-binding proteins: the *S. cerevisiae* DEAD-box ATPases Dhh1 and Ded1, and the *C. elegans* P-granule protein PGL-3. We tested five factors known to affect condensation: ATP (0 – 4 mM), NaCl (100 – 200 mM), synthetic RNA homopolymer poly(U) (0 – 100 µg/mL), pH (6.5 – 7.5) and protein concentration (5 – 10 µM).

To identify the factors with the strongest effects, we first used a definitive screening design (DSD). For the five factors tested here, the DSD required 11 conditions ((2m+1), where m denotes the number of factors tested)^41,42^. Across these conditions, all five factors were varied at defined low, intermediate or high levels (Supplementary Table 1a). This allowed us to estimate the ‘main effect’ of each factor across the tested design space. By comparison, a two-level full factorial design that tests all possible combinations would require 2⁵ = 32 conditions per protein. At this screening stage, the analysis focused on the main effect of each factor and did not yet consider pairwise interactions between factors or nonlinear responses.

Condensation was quantified from fluorescence images using the coefficient of variation (CV) of pixel intensities. CV increases as protein redistributes from a comparatively uniform diffuse signal into bright condensate assemblies (Supplementary Fig. 1).

To enhance robustness of the results, the 11-condition DSD was performed in two independent ‘experimental blocks’ (replicates), each using an independently purified and labelled protein preparation. Data from both blocks were analysed together in a single regression model, with experimental block included as another ‘variable’ alongside ATP, pH, poly(U), protein concentration and NaCl. The experimental block variable accounts for systematic differences between the two independent experiments, allowing the effects of the five experimental factors to be estimated independently of block-to-block variation.

The models estimate the magnitude and direction of the effect of each factor on condensation. A positive effect indicates that increasing the level of a factor increases CV and thus promotes condensation, whereas a negative effect indicates that increasing the factor decreases CV and thus reduces condensation. To compare the five factors on a common scale, each estimated effect was divided by its standard error. These standardised effects are displayed as Pareto plots, which rank the factors according to the magnitude of their effect on condensates (Fig. 1d–f). The α = 0.05 threshold indicates which effects reached statistical significance. Because these models include only the main effect of each factor, they do not capture pairwise interactions between factors or nonlinear responses.

The screening models showed moderate overall fit (R²) but limited predictive performance (predicted R²), particularly for Dhh1 and Ded1 (Supplementary Table 1c). PGL-3 and Ded1 showed differences in overall response between the two experimental blocks. Nevertheless, separate analysis of the two experimental blocks showed that the directions and magnitudes of the main effects of pH, poly(U), protein-concentration, ATP and NaCl were comparable (Supplementary Fig. 2), indicating that the principal trends identified by the screen are robust to block-to-block variation.

At this exploratory screening stage, several trends were consistent across all three proteins: lower NaCl and higher protein concentration generally favoured condensation, whereas ATP produced only small effects and did not promote condensation within the tested range. This was unexpected for Dhh1 because nucleotide binding is normally closely linked to RNA binding and conformational cycling in DEAD-box ATPases^16,37,43–45^.

Protein-specific responses that exceeded the α = 0.05 significance threshold differed most strongly for pH and poly(U). For Dhh1, lower pH and poly(U) both promoted condensation (Fig. 1d). Ded1 also showed a significant pH effect, with lower pH favouring condensation, whereas poly(U) showed an inhibitory trend (Fig. 1e). For PGL-3, poly(U) significantly promoted condensation, while pH had a much smaller effect (Fig. 1f). The remaining effects were interpreted as preliminary trends at this screening stage. Thus, the same biochemical inputs produce distinct responses depending on the protein, suggesting protein-specific regulatory logic in how these inputs are integrated.

The limited predictive performance of the screening results indicate that main factor effects alone are insufficient to describe the full condensation behaviour of the three proteins. We therefore reduced the design space for more detailed analysis.

The screening models showed that pH and the concentrations of poly(U) and NaCl produced clear and protein-specific responses, making them the most interesting factors for more detailed analysis. We therefore carried these three factors forward into the subsequent full-factorial design. Protein concentration was fixed at 5 µM to reduce protein consumption and avoid extensive condensation.

### Factorial analysis resolves pairwise interactions

In the next step, we asked whether the factors prioritized by the DSD screen - pH, poly(U) and NaCl - act independently or ‘interact’ with each other, meaning that the effect of one factor depends on the level of another factor. To this goal, we performed a 2-level full factorial experiment with centre points. Each factor was tested at a low and a high level. The resulting 2^3^ factorial design contained all eight possible combinations of conditions, corresponding to the corners of a cube in the three-dimensional design space (Fig. 2a). This design allowed us to estimate both the main effect of each factor and pairwise interactions between factors.

**Fig. 2.**
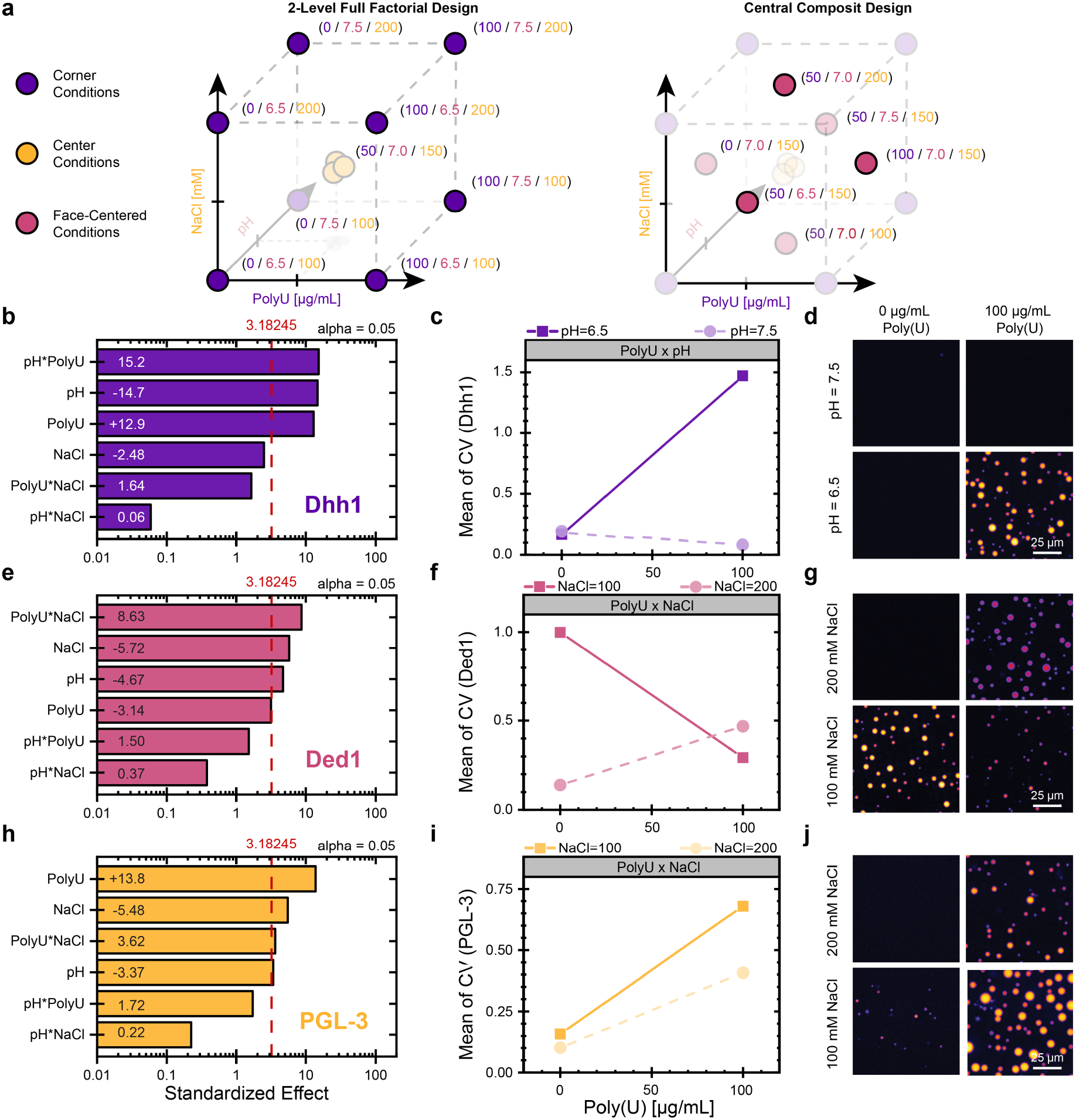
| Full-factorial analysis reveals protein-specific interactions between pH, RNA and ionic strength. **a** Schematic representation of the 2-level full-factorial design (left) and the central composite face-centred (CCF) design (right). The axes represent poly(U) concentration, pH and NaCl concentration in the design space. Point labels indicate design space coordinates in the following order (poly(U) / pH / NaCl). Purple dots represent the eight corner conditions sampled by the 2-level full-factorial design and the yellow dots represent the centre condition. In the CCF schematic, pale purple dots indicate conditions inherited from the full-factorial design and red dots indicate the additional face-centred conditions used for response-surface modelling. **b–d** Dhh1 full-factorial analysis: **b** Pareto plot of standardised main and interaction effects. **c** Interaction plot showing the dominant pH–poly(U) interaction. Poly(U) strongly promotes condensation at low pH (solid line: pH 6.5) but has little effect at high pH (dashed line: pH 7.5). **d** Representative microscopy images of the corresponding conditions. **e–g** Ded1 full-factorial analysis: **e** Pareto plot of standardised main and interaction effects. **f** Interaction plot showing the dominant poly(U)–NaCl interaction. Poly(U) inhibits condensation at low NaCl (solid line: 100 mM) but promotes condensation at high NaCl (dashed line: 200 mM). **g** Representative microscopy images of the corresponding conditions. **h–j** PGL-3 full-factorial analysis: **h** Pareto plot of standardised main and interaction effects. **i** Interaction plot showing the dominant poly(U)–NaCl interaction. Poly(U) promotes condensation most strongly at low NaCl (solid line: 100 mM), with a weaker response at high NaCl (dashed line: 200 mM). **j** Representative microscopy images of the corresponding conditions. Condensation was quantified using the coefficient of variation (CV) of the protein fluorescence signal, and the red dashed line marks the α = 0.05 significance threshold. Scale bars, 25 µm.

We also included a centre-point condition in which all three factors were set to their intermediate levels, which was measured in three independently prepared ‘runs’ (reactions). These replicated centre points provided an estimate of experimental variability and allowed us to test for ‘curvature’, which occurs when the response at the centre of the design space deviates from that predicted by a linear model. Significant curvature (p < 0.05) therefore indicates that the response cannot be fully described by linear effects. In total, the experiment contained nine distinct conditions measured in eleven runs per protein: eight factorial corner point and three centre point runs (Supplementary Table 2).

The measured CV values were fitted with first-order models containing three main effects and three pairwise interaction terms. The models closely fitted the measured data, with R² values of 0.997 for Dhh1, 0.979 for Ded1 and 0.991 for PGL-3. Predicted R² was high for Dhh1 (0.954) and Ded1 (0.928) but lower for PGL-3 (0.472).

The full-factorial measurements confirmed the main trends identified in the DSD screen and revealed interactions that were not captured by the DSD screening models. For Dhh1, low pH and high poly(U) concentration promoted condensation significantly, whereas high NaCl concentration had only a weak negative effect (Fig. 2b). The strongest interaction occurred between pH and poly(U): poly(U) promoted condensate formation at pH 6.5 but was insufficient to support robust condensation at pH 7.5 (Fig. 2c, d).

To test whether pH-dependent RNA binding explains this factor interaction, we measured the RNA-binding affinity of Dhh1 using fluorescence anisotropy but could not detect a significant change in binding affinity across the tested pH range (Supplementary Fig. 3). The pH dependence of Dhh1 condensation therefore cannot be explained by changes in RNA affinity alone and may instead arise from other pH-sensitive contributions to condensate formation, such as Dhh1 self-interactions.

Ded1 showed a different response compared to Dhh1. Low pH and low NaCl favoured condensation, and the strongest interaction occurred between poly(U) and NaCl (Fig. 2e). At 100 mM NaCl, Ded1 formed condensates without poly(U), and addition of poly(U) reduced condensation. At 200 mM NaCl, condensation was strongly reduced in the absence of poly(U) but partially restored in the presence of poly(U) (Fig. 2f, g). The effect of poly(U) on Ded1 condensation therefore depends on ionic strength, switching from inhibitory at low NaCl to promoting at high NaCl.

PGL3 showed a third, distinct regulatory pattern. Poly(U) was the strongest positive factor, whereas increasing NaCl and pH reduced condensation (Fig. 2h). The strongest pairwise interaction was again between poly(U) and NaCl, but in contrast to Ded1, poly(U) promoted PGL-3 condensation most strongly at low ionic strength and at low NaCl concentrations PGL-3 condensed even in the absence of poly(U) (Fig. 2i, j).

The factorial analysis therefore identifies pH, RNA and ionic strength as shared influential factors, but shows that their individual main effects and pairwise interactions differ between proteins. There was no evidence of curvature for Ded1 (p = 0.85), indicating that its response was well described by the linear model. By contrast, significant curvature was detected for Dhh1 (p = 4.1 × 10⁻⁴) and PGL-3 (p = 3.6 × 10⁻³).

The full-factorial models thus resolve important pairwise interactions, but the significant curvature observed for Dhh1 and PGL-3 indicates that linear models are not sufficient to fully capture their responses. We therefore expanded the experimental design in the next step to map responses across the multifactorial space between the tested conditions.

### Response surfaces map condensation landscapes

The two-level full factorial experiments sampled only the eight corners and replicated centre of the selected design space. To determine how condensation varied between the discrete conditions, we conducted a face-centred central composite design (CCF) as a separate experiment. All conditions corresponding to the original two-level full-factorial design were independently prepared and measured again. In addition, the CCF included six new face-centre conditions, with one located at the centre of each cube face (Fig. 2a).

The resulting dataset comprised eight corner conditions, six face-centre conditions and six independently prepared centre-point runs. This increased coverage allowed us to detect nonlinear responses and construct continuous response surfaces across the investigated design space. The complete design matrices and factor levels are provided in Supplementary Table 3.

We first fitted full quadratic models containing three linear terms, three pairwise interaction terms and one quadratic term for each factor. These terms represent the main effect of each factor (linear main effect terms), whether the effect of one factor depends on the level of another (pairwise interaction terms), and the curvature in the response (quadratic terms).

Although the full models fitted the measured data well, their predictive performance was limited (Supplementary Table 3c). This indicated ‘overfitting’, meaning that some model terms captured experimental variation rather than reproducible response patterns. Overfitting was particularly pronounced for Dhh1 and Ded1, whose adjusted R² values (0.71 and 0.64) contrasted with negative predicted R^2^ values (-0.30 and -0.24). PGL-3 showed a milder degree of overfitting, with an adjusted R² of 0.91 and a predicted R² of 0.50 (Supplementary Table 3c).

We therefore simplified the model: all three main factors were retained, but only significant pairwise interaction and quadratic terms were included (Fig. 3a-c). This substantially improved predictive performance for all three proteins (-0.30 → 0.65 for Dhh1, -0.24 → 0.64 for Ded1 and 0.5 → 0.76 for PGL3). The reduced CCF models therefore provide a better balance between describing the measured response and avoiding overfitting.

**Fig. 3.**
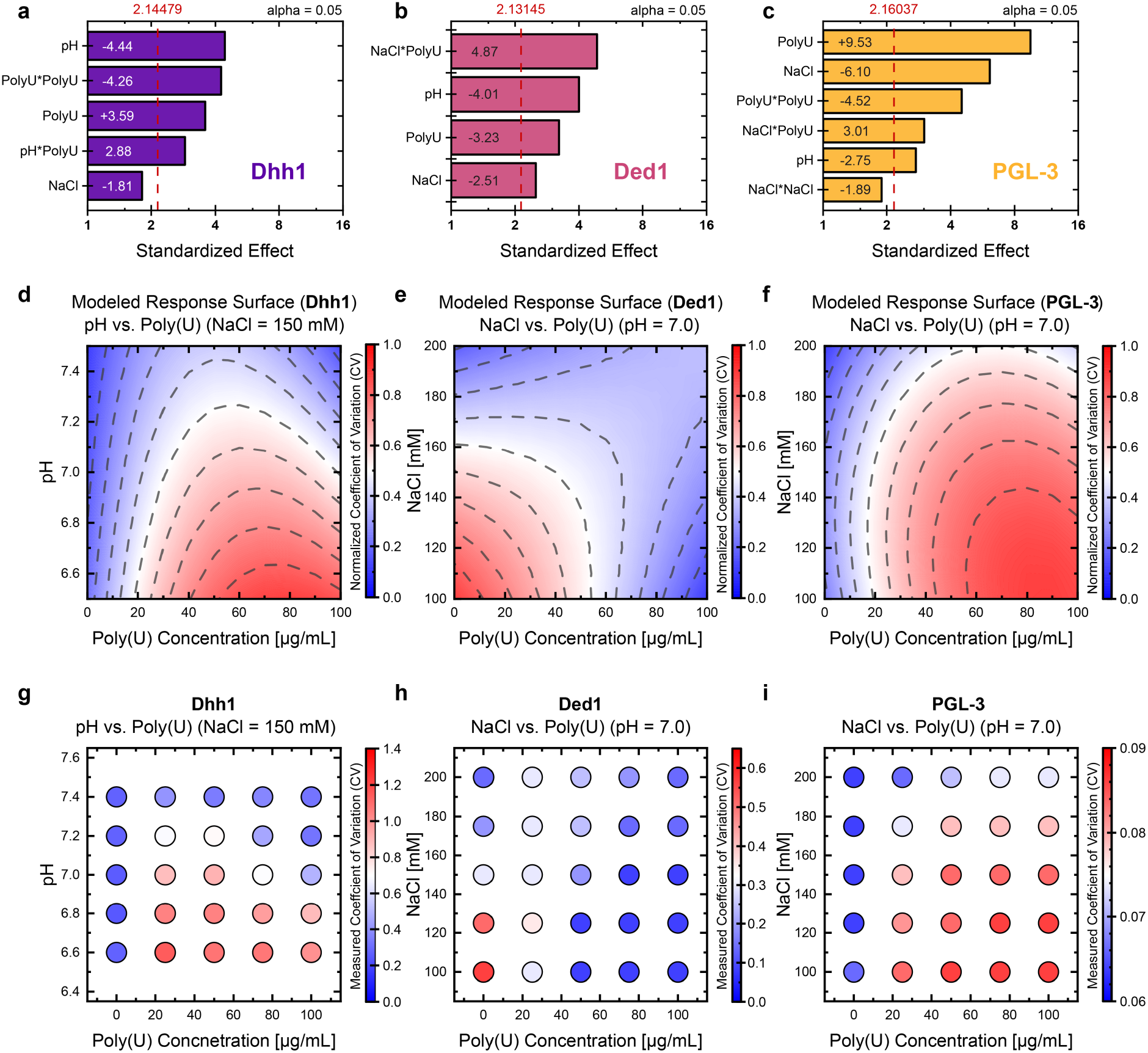
| Response-surface modelling captures protein-specific nonlinear condensation landscapes. **a–c** Pareto plots from the reduced central composite face-centred (CCF) models for Dhh1, Ded1 and PGL-3. Condensation was quantified using the coefficient of variation (CV) of the protein fluorescence signal and bars show the standardised effects retained in the reduced models, including main effects, pairwise interactions and quadratic terms. The red dashed line marks the α = 0.05 significance threshold. **d–f** Normalised response surfaces predicted from the reduced CCF models. The surfaces show the expected condensation response across the selected two-factor design spaces. For Dhh1, the surface maps pH and poly(U) concentration. For Ded1 and PGL-3, the surfaces map NaCl and poly(U) concentration. Red regions indicate higher predicted CV values and blue regions indicate lower predicted CV values. **g–i** Independent benchmarking measurements across the same two-factor spaces. These data are shown as measured non-normalised CV values and are therefore displayed separately from the normalised model surfaces. Benchmarking was performed in three independent experiments and recapitulated the main trends predicted by the CCF models (n = 3).

The resulting response surfaces extend the pairwise interaction patterns identified by the factorial experiments across continuous representations of the investigated design space. For Dhh1, condensation is strongest at low pH in the presence of poly(U), consistent with the pH– poly(U) interaction identified in the factorial analysis (Fig. 3d). The Ded1 surface captures the context-dependent effect of poly(U). RNA suppresses condensation at low NaCl but promotes condensation as NaCl increases (Fig. 3e). PGL-3 shows an opposite RNA–salt relationship, with poly(U) promoting condensation most strongly at low NaCl (Fig. 3f). The same biochemical variables therefore produce substantially different response landscapes for each of the three proteins.

Significant lack-of-fit nevertheless remained for Dhh1 and PGL-3, indicating that some local response variation was not captured by the quadratic model. The resulting response surfaces should therefore be interpreted as representations of the main trends across the investigated design space rather than precise predictions for every individual condition.

To test whether the modelled trends reflected experimental behaviour, we performed independent benchmarking measurements for two factors defining the selected response surface (Fig. 3d-f). The measurements are shown separately because the response surfaces display normalised predicted values, whereas the benchmarking experiments report non-normalised CV values (Fig. 3g-i). For all three proteins, the independent benchmark measurements reproduced the main trends predicted by the CCF models. Across 25 benchmarking conditions per protein, predicted responses correlated positively with mean measured CV, with Pearson’s r values of 0.865 for Dhh1, 0.734 for Ded1 and 0.956 for PGL-3 (Supplementary Fig. 4).

The CCF analysis thus extends the discrete interaction patterns into continuous descriptions of the investigated design space. The resulting response surfaces reveal distinct condensation landscapes for Dhh1, Ded1 and PGL-3 and provide experimentally benchmarked guides to their behaviour across the tested conditions.

### Protein mixtures form distinct internal architectures

Having mapped the condensation landscapes of the individual proteins, we next asked how the presence of a second protein influenced these responses. We performed separate two-level factorial co-condensation experiments for each protein pair, varying pH, NaCl, poly(U) concentration and the presence or absence of the ‘secondary’ protein. Condensation of the ‘primary’ protein of interest was quantified using its CV, and each experiment was performed in three independent experimental blocks (replicates). Consequently, the effect magnitudes in these models reflect responses within the respective experimental framework and are not expected to be identical between protein-pair experiments or to those obtained in the earlier single-protein factorial analysis. The resulting full-factorial models showed good predictive performance, with predicted R² values of 0.73 for PGL3-Dhh1, 0.70 for PGL3-Ded1 and 0.92 for Dhh1-Ded1 (Supplementary Table 4).

In mixtures containing PGL-3 as the primary protein, addition of either Dhh1 or Ded1 had only a small effect on the PGL-3 condensation response (Fig. 4a, b). The main effect of the partner protein and pairwise interaction terms involving the partner protein were small relative to the effects of pH, NaCl and poly(U), indicating that neither Dhh1 nor Ded1 substantially altered how PGL-3 responded to the biochemical conditions within the respective experiments.

**Fig. 4.**
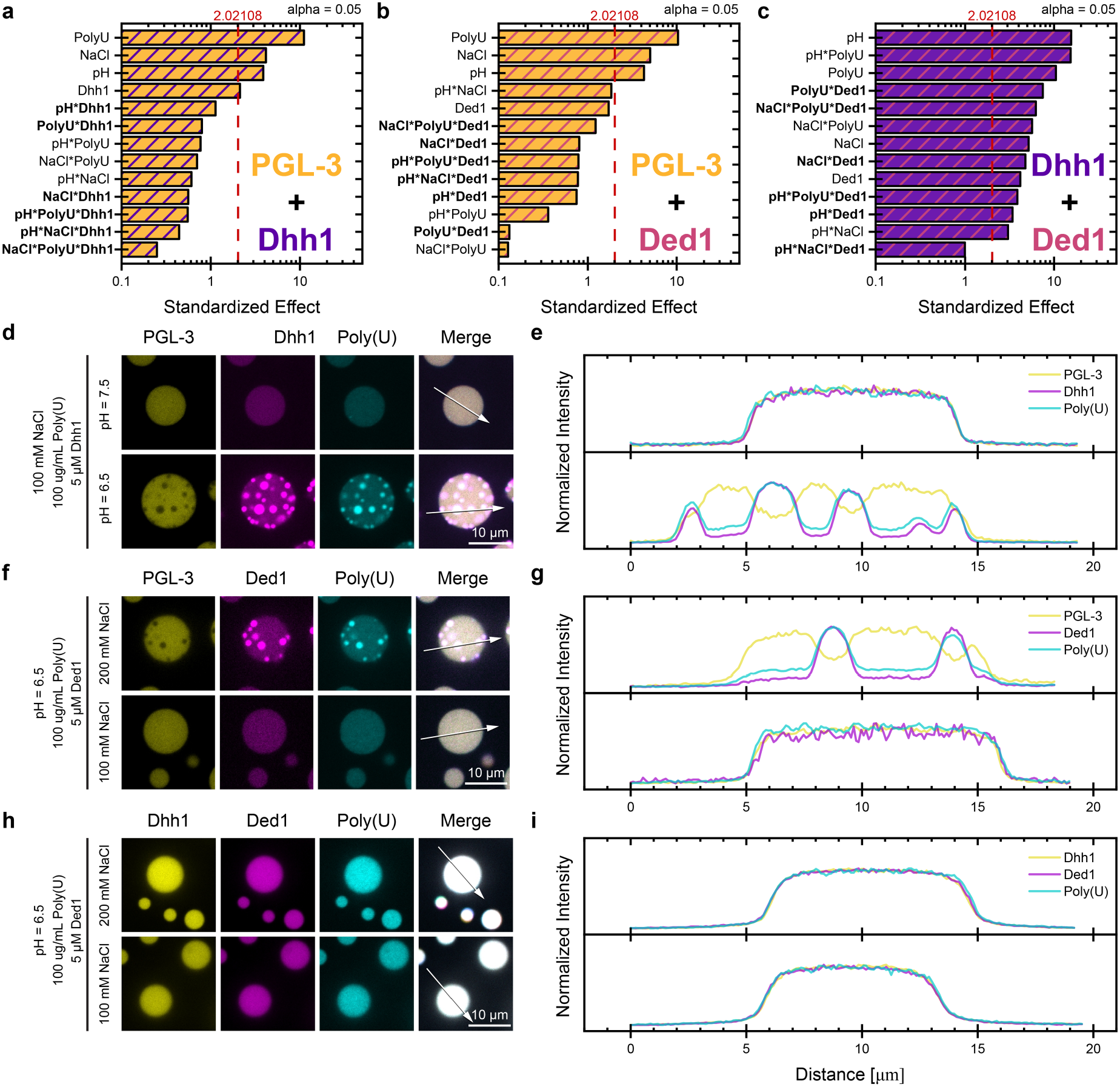
| Co-condensation effects and internal organisation depend on the protein pair. **a–c** Pareto plots from two-level factorial co-condensation experiments for **a** PGL-3–Dhh1, **b** PGL-3–Ded1 and **c** Dhh1–Ded1 mixtures. The response was quantified using the coefficient of variation (CV) of the primary protein fluorescence signal. Bars represent the standardised main and interaction effects. Interaction terms involving the secondary protein are highlighted in bold. The red dashed line marks the α = 0.05 significance threshold. PGL-3 was used as the primary protein in **a, b** and Dhh1 in **c**. Each experiment was performed in three independent experimental blocks. **d, e** PGL-3–Dhh1 co-condensates. **d** Representative condensates at pH 7.5 and pH 6.5. **e** Corresponding normalised fluorescence intensity profiles along the indicated line scans. Dhh1 and poly(U) form internal substructures within PGL-3 condensates at pH 6.5 but remain more uniformly distributed at pH 7.5. **f, g** PGL-3–Ded1 co-condensates. **f** Representative condensates at 200 and 100 mM NaCl. **g** Corresponding normalised fluorescence intensity profiles. Ded1 and poly(U) form internal substructures at 200 mM NaCl but are more uniformly distributed at 100 mM NaCl. **h, i** Dhh1–Ded1 co-condensates. **h** Representative condensates at 200 and 100 mM NaCl. **i** Corresponding normalised fluorescence intensity profiles showing overlapping Dhh1, Ded1 and poly(U) signals without internal demixing. Scale bars, 10 µm.

The internal organisation of the mixed condensates, however, depended strongly on the condensation behaviour of the partner proteins. In PGL3-Dhh1 mixtures, Dhh1 partitioned weakly and relatively uniformly into PGL-3 droplets under conditions that did not support condensation of Dhh1 itself on its own. By contrast, under conditions that favoured Dhh1 condensation in the single-protein experiments, most notably low pH in the presence of poly(U), Dhh1 formed discrete internal condensate-within-condensate structures that were depleted of PGL-3 and enriched in poly(U) (Fig. 4d, e). Ded1 showed a similar behaviour: under conditions that favoured Ded1 condensation in presence of poly(U), particularly low pH and high NaCl, it formed RNA-rich condensates-within-condensates inside PGL-3 droplets (Fig. 4f, g). The single-protein landscapes therefore not only report whether Dhh1 and Ded1 condense but also indicate when they reorganise into internal substructures within PGL-3 condensates.

A different behaviour emerged when Dhh1 and Ded1 were combined. Here, Ded1 measurably altered the Dhh1 condensation landscape, contributing several significant interaction terms to the model. The model retained high predictive performance (predicted R² = 0.92; Fig. 4c). Unlike PGL-3 mixtures, Dhh1-Ded1 mixtures did not form obvious condensate-within-condensate structures. Instead, Dhh1, Ded1 and poly(U) remained relatively uniformly distributed across the co-condensate (Fig. 4h-i).

These co-condensation experiments show that changes in the condensation response of the primary protein and changes in the internal organisation of co-condensates are distinct properties of protein mixtures. Significant pairwise interactions involving the secondary protein indicate that the effect of a biochemical factor (e.g. salt, pH) on the primary protein depends on the presence of the secondary protein. By contrast, single-protein landscapes can indicate conditions in which a recruited protein is more likely to form internal substructures. However, neither is sufficient to predict co-condensate organisation. Co-condensation therefore depends on the specific protein combination and ultimately needs to be examined directly.

### Dhh1 condenses with RNA without added ATP

Having established that the sequential DoE workflow captured reproducible, context-dependent condensation behaviour, we next asked whether unexpected responses emerging from the screen could identify molecular determinants of condensate assembly. We focused on Dhh1, where ATP failed to promote condensation in the initial screen (Fig. 1d) despite the established coupling between nucleotide state and canonical RNA engagement in DEAD-box ATPases^16,37,43–45^.

This prompted us to examine Dhh1 condensation across ATP concentrations directly. Across a targeted ATP titration, Dhh1 readily formed condensates with poly(U) RNA in the absence of ATP (Supplementary Fig. 5) and at low ATP concentrations (Fig. 5a). Increasing ATP progressively reduced condensation, with an apparent IC₅₀ of 3.8 ± 0.4 mM (Fig. 5b), which goes hand-in-hand with the suggested role of ATP as a biological hydrotrope^15^. ATP is therefore not required for Dhh1 condensation under these assay conditions and suppresses condensation at higher concentrations.

**Fig. 5.**
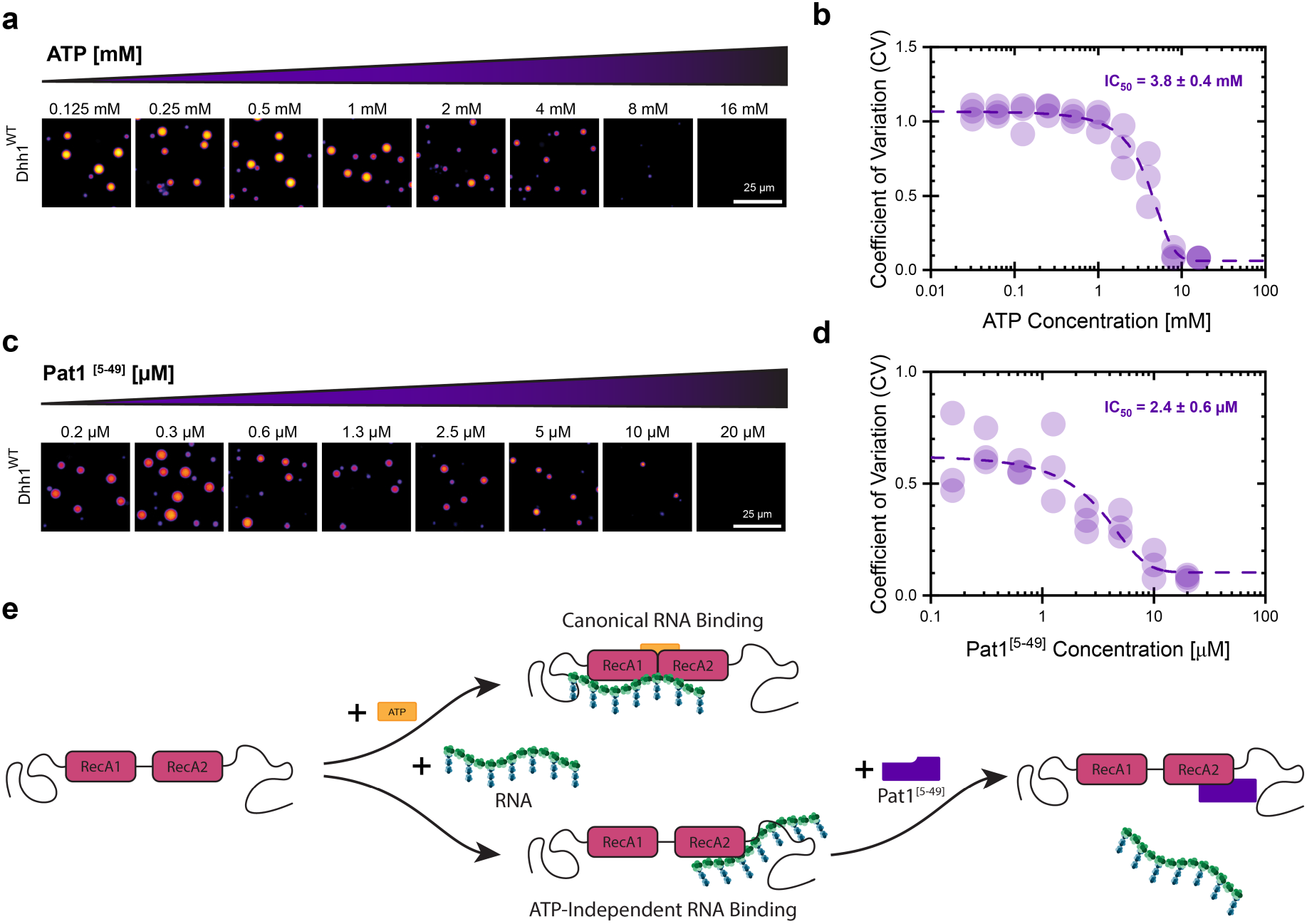
|Dhh1 forms condensates independently of added ATP. **a** Representative fluorescence microscopy images of Dhh1–poly(U) condensates across an ATP titration. Assay conditions: pH = 7.0, 50 µg/mL poly(U) and 150 mM NaCl **b** Quantification of Dhh1 condensation shown in **a** using the coefficient of variation (CV) of the Dhh1 fluorescence signal, resulting in an apparent IC₅₀ of 3.8 ± 0.4 mM. (n = 3) **c** Representative fluorescence microscopy images of Dhh1–poly(U) condensates across a Pat1^[5–49]^ titration. Assay conditions: pH = 7.0, 50 µg/mL Poly(U), 150 mM NaCl and no added ATP **d** CV-based quantification Dhh1 condensation shown in **c** with an apparent IC₅₀ of 2.4 ± 0.6 µM. (n = 3) **e** Working model of the two RNA-binding modes of Dhh1. RNA can engage the canonical ATP-dependent RNA-binding site or a secondary ATP-independent RNA-binding surface on the RecA2 domain. Pat1^[5–49]^ binds to an overlapping region on the RecA2 domain and inhibits ATP-independent Dhh1 condensation. Scale bars, 25 µm.

The ability of poly(U) to support condensation without ATP suggested that ATP-independent Dhh1–RNA interactions could be important for condensate formation. Structural studies previously identified a noncanonical, ATP-independent RNA-binding surface on the RecA2 domain of Dhh1 that overlaps with the binding site for the N-terminal region of Pat1, another P-body protein^46^. Furthermore, the N-terminal fragment of Pat1 was previously shown to suppress Dhh1 condensation in the presence of ATP^47^.

We therefore tested whether the N-terminal region of Pat1 (Pat1^[5–49]^) also affected ATP-independent Dhh1–poly(U) condensation. Increasing concentrations of Pat1^[5–49]^ progressively reduced condensation, with an apparent IC₅₀ of 2.4 ± 0.6 µM (Fig. 5c, d), indicating that this secondary RNA-binding surface is indeed important for ATP-independent Dhh1 condensation.

This targeted analysis confirms the unexpected observation from the DoE screen: Dhh1 does not require its canonical ATP-dependent RNA-binding mode to form condensate. Instead, ATP-independent RNA binding involving a secondary surface on RecA2 can support Dhh1 condensation. These findings are consistent with a model in which Dhh1 interacts with RNA through two modes: canonical ATP-dependent RNA binding and ATP-independent RNA binding through the secondary RecA2 surface (Fig. 5e). Pat1^[5–49]^ binds to an overlapping site on RecA2 and inhibits ATP-independent Dhh1 condensation, consistent with competition at this secondary RNA-binding surface (Fig. 5e). The RecA2 RNA-binding surface promotes efficient Dhh1 RNA binding and condensation.

To test directly whether the secondary RNA-binding surface contributes to Dhh1 condensation, we used two previously characterized mutants in the RecA2 domain: Dhh1^Mut1^ (N269A, H272A, T276A & K280A) and Dhh1^Mut3^ (S292D, N294D & R295D) (Fig. 6a)^46^. Both mutants were originally studied for their effects on the Dhh1-Pat1 interaction: Mut1 reduced Pat1 binding, whereas Mut3 was reported to completely abolish the interaction.

**Fig. 6.**
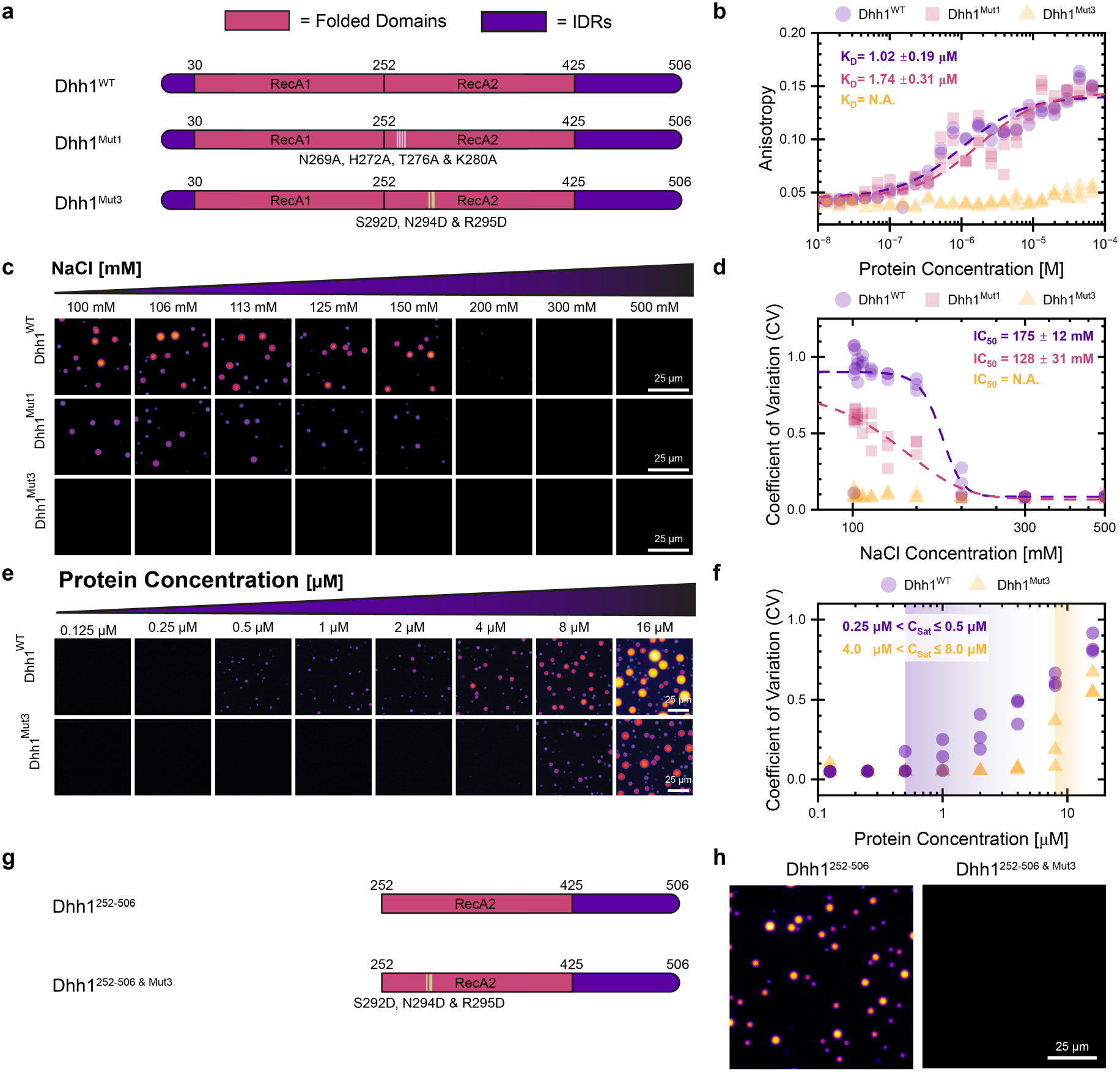
| Mutations in the secondary RNA-binding surface on the RecA2 domain impair RNA binding and condensation. **a** Domain organisation of Dhh1^WT^ and the Dhh1^Mut1^ (N269A, H272A, T276A & K280A) and Dhh1^Mut3^ (S292D, N294D & R295D) variants carrying mutations within the secondary RNA-binding surface of the RecA2 domain. Folded RecA domains and intrinsically disordered regions are indicated. **b** Fluorescence anisotropy measurements of Dhh1ᵂᵀ Dhh1ᴹᵘᵗ¹ and Dhh1ᴹᵘᵗ³ binding to an 18-mer oligo(U) RNA. (n = 3) **c, d** NaCl titration experiment comparing condensation of Dhh1^WT^ Dhh1^Mut1^ and Dhh1^Mut3^ at 5 µM protein concentration. Assay Conditions: pH = 7.0, 50 µg/mL poly(U) and no added ATP (n = 3). **c** Representative fluorescence microscopy images. **d** CV-based quantification of the observed condensation. Dhh1^Mut1^ showed increased salt sensitivity relative to Dhh1ᵂᵀ whereas Dhh1^Mut3^ did not form detectable condensates across the tested NaCl range. **e, f** Protein-concentration titration experiment comparing condensation of full-length Dhh1ᵂᵀ and Dhh1^Mut3^. Assay Conditions: pH = 7.0, 50 µg/mL poly(U), 100 mM NaCl and no added ATP (n = 3). **e** Representative fluorescence microscopy images. **f** CV-based quantification showing that higher protein concentrations partially restored condensation of Dhh1^Mut3^. **g** Domain organization of the C-terminal Dhh1^252–506^ truncation and the corresponding Dhh1^252–506^ ^+ Mut3^ variant. **h** Representative fluorescence microscopy images of Dhh1^252–506^ and Dhh1^252–506^ ^+ Mut3^ at 16 µM protein concentration showing condensation of the unmutated truncation but not the Mut3 variant. Assay Conditions: pH 7.0, 50 µg/mL poly(U) and 100 mM NaCl and no added ATP (n = 3). Scale bars, 25 µm.

We first tested whether the introduced mutations also affect RNA binding. Fluorescence anisotropy showed that wild-type Dhh1 and Dhh1^Mut1^ bound an U_18_ RNA oligonucleotide with similar apparent affinity (K_D_ = 1.02 ± 0.19 µM resp. 1.74 ± 0.31 µM). By contrast, RNA binding of Dhh1^Mut3^ was strongly impaired and could not reliably be fitted to a binding isotherm (Fig. 6b).

If the secondary ATP-independent RNA binding surface promotes condensation, mutations that perturb this region should reduce Dhh1 condensation. We therefore compared Dhh1^WT^ with the Mut1 and Mut3 variants. To increase the sensitivity of this comparison, we measured condensation across a range of NaCl concentrations. Wild-type Dhh1 formed condensates at low NaCl concentrations and progressively dissolved as NaCl concentration increased, with an apparent IC_50_ of 175 ± 12 mM NaCl (Fig. 6c, d). Dhh1^Mut1^ remained condensation-competent but was more sensitive to salt, with an apparent IC₅₀ of 128 ± 31 mM. This partial reduction in condensation is consistent with Mut1 weakening, rather than completely disrupting, the secondary interaction surface.

Dhh1^Mut3^ showed a much stronger phenotype and did not form detectable condensates under all tested conditions. Increasing the protein concentration partially rescued condensation: wild-type Dhh1 had an apparent saturation concentration (c_sat_) of approximately 0.5 µM at 100 mM NaCl, whereas Dhh1^Mut3^ required substantially higher protein concentrations, with an apparent c_sat_ of approximately 8 µM (Fig. 6e, f). Mut3 therefore strongly reduces the ability of Dhh1 to condense but does not abolish it completely.

We next asked whether the C-terminal region containing the secondary RNA binding site is sufficient to support condensation and generated a Dhh1 construct (Dhh1^252-506^) containing RecA2 and the C-terminal intrinsically disordered region (IDR) (Fig. 6g). This fragment remained condensation-competent and started to form condensates at 16 µM protein (Fig. 6h). Introducing the Mut3 substitutions into this construct abolished detectable condensation. These results show that Dhh1^252-506^ can support condensation independently of the N-terminal half of Dhh1 and that the Mut3-sensitive surface is important for this ability.

The mutational and truncation analyses therefore support a model in which the secondary, ATP-independent RecA2 RNA-binding surface, together with the C-terminal IDR, makes a major contribution to condensation. However, the isolated C-terminal half (Dhh1^252-506^) has an approximately 30-fold higher apparent saturation concentration than the full-length protein, indicating that additional regions also contribute to efficient condensation.

### The RecA2 surface supports P body formation

Having shown that the Mut3 substitutions impair RNA binding and Dhh1 condensation *in vitro*, we next asked whether they also affect Dhh1-dependent P-body formation in yeast. Previous work showed that Dhh1^Mut3^ fails to support efficient P-body assembly, a phenotype attributed primarily to the disruption of the Dhh1–Pat1 interaction^48^. Because our biochemical experiments additionally showed that Mut3 strongly impairs RNA binding and condensation, we re-examined the contribution of this region to P-body formation in cells lacking endogenous Dhh1. P-body assembly was quantified using endogenously mCherry-tagged Dcp2, as a P-body marker. Following glucose depletion, control cells expressing endogenously yeGFP-tagged Dhh1 formed 0.67 ± 0.24 Dcp2-mCherry positive P-bodies per cell. By contrast, cells lacking endogenous Dhh1 (*dhh1Δ*) formed significantly fewer P-bodies per cell (0.06 ± 0.01 per cell), consistent with the established role of Dhh1 in P-body assembly^37^ (Fig. 7a,c).

**Fig. 7.**
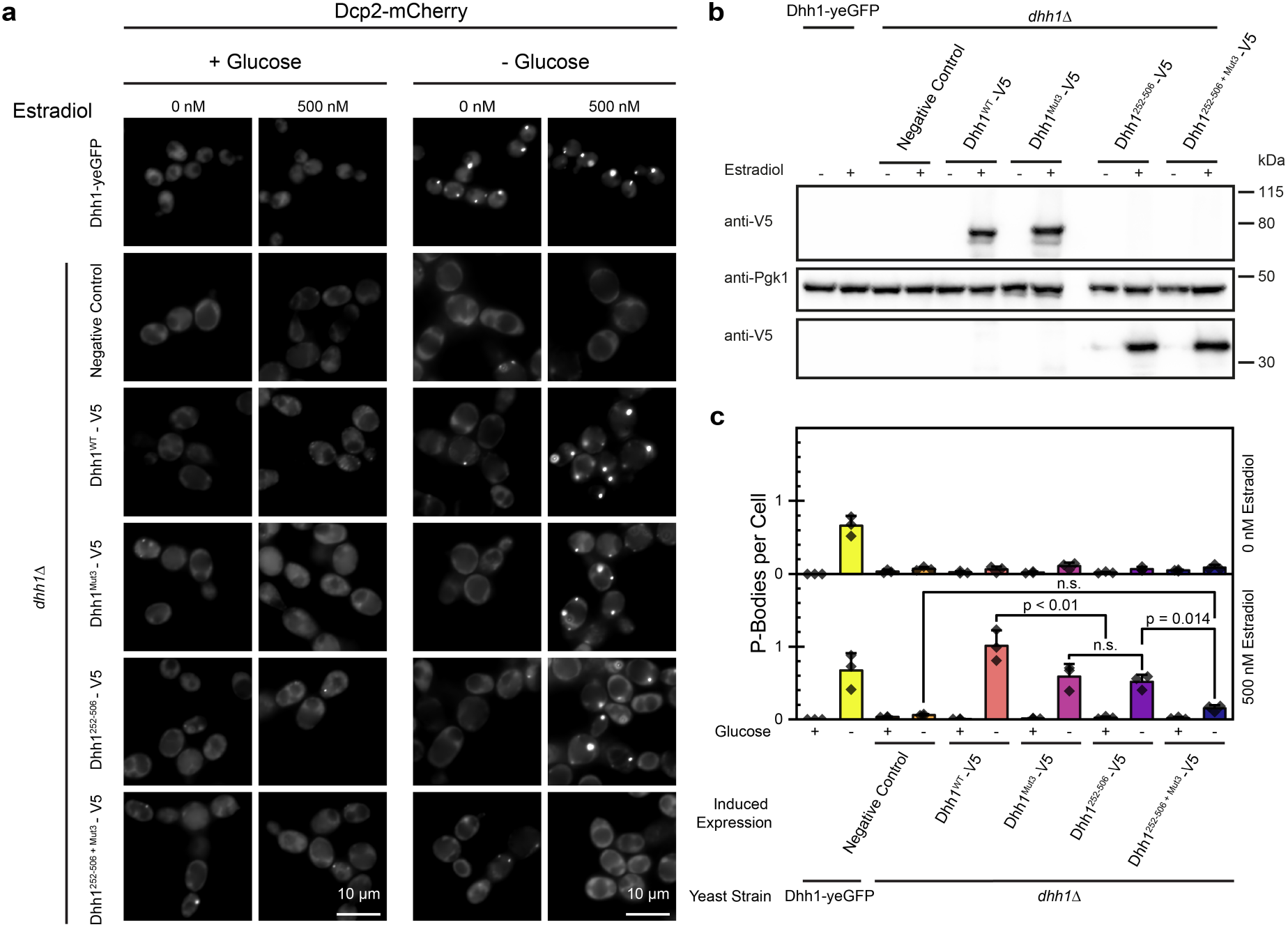
| The Mut3-sensitive RecA2 surface supports Dhh1-dependent P-body formation in yeast. **a** Representative fluorescence microscopy images of Dcp2-mCherry-positive P-bodies under normal growth conditions and following glucose depletion. Endogenous Dhh1-YEGFP and *dhh1Δ* cells are shown alongside *dhh1Δ* cells carrying estradiol-inducible Dhh1^WT^–V5, Dhh1^Mut3^–V5, Dhh1^252-506^–V5 or Dhh1^252-506^ ^+ Mut3^–V5 constructs in the absence or presence of 500 nM estradiol. Images were denoised using Nikon Denoise.ai. Scale bars, 10 µm. **b** Western Blot of the corresponding estradiol-inducible Dhh1 constructs. Anti-V5 immunoblotting confirmed estradiol-dependent expression of the full-length and truncated Dhh1 constructs at their expected molecular masses. Pgk1 was detected as a loading control. **c** Quantification of Dcp2-positive P-bodies per cell under the corresponding conditions shown in **a**. Data represent mean ± standard deviation from three independent biological replicates, with ≥150 cells quantified per replicate (n = 3). Statistical comparisons were corrected for multiple testing using the Holm–Bonferroni method; significance threshold, p < 0.05.

To compare the effect of different Dhh1 variants, we expressed V5-tagged constructs in *dhh1Δ* cells using an estradiol-inducible LexA system^49^. In the absence of estradiol and thus construct expression, none of the proteins was expressed, and none of the strains showed P-body formation in glucose-depleted cells. Following induction with 500 nM estradiol, all V5-tagged constructs were expressed at their expected molecular masses (Fig. 7b). Full-length Dhh1ᵂᵀ-V5 restored robust P-body formation (1.01 ± 0.21 P-bodies per cell) (Fig. 7a, c). The endogenous reference strain expressed Dhh1–yeGFP, whereas the complementation constructs carried a V5 tag and were expressed from an inducible promoter. Because differences in tag and expression context could affect P-body formation^50^, we did not directly compare absolute P-body numbers between the endogenous yeGFP-tagged reference strain and the V5-tagged, inducible complementation constructs. Instead, the mutant and truncation constructs were compared with Dhh1^WT^–V5 expressed under the same induction conditions.

The mutant of the RecA2 secondary RNA binding surface (Dhh1^Mut3^–V5) produced 0.59 ± 0.17 P-bodies per cell, fewer than Dhh1^WT^–V5 control but significantly more than the *dhh1Δ* control. The Mut3-sensitive RecA2 surface therefore contributes to efficient P-body assembly but is not strictly required in the context of full-length Dhh1.

We next tested the C-terminal construct (Dhh1^252-506^–V5), which lacks the RecA1 domain and the N-terminal IDR (Fig. 6g). Cells expressing Dhh1^252-506^–V5 formed 0.52 ± 0.10 P-bodies per cell, similar to full-length Dhh1^Mut3^–V5. Introducing the Mut3 substitutions into this truncated construct strongly reduced P-body formation: cells expressing Dhh1^252-506+Mut3^–V5 formed only 0.15 ± 0.04 P-bodies per cell. The C-terminal half of Dhh1 by itself can therefore support P-body formation in cells, but the Mut3 substitutions strongly impair this ability. The weaker effect of Mut3 in full-length Dhh1 compared to the truncated construct suggests that additional regions can partially compensate when this RecA2 region is perturbed.

Together, the cellular experiments are consistent with previous evidence that the RecA2 region affected by Mut3 contributes to P-body assembly^48^. Importantly, our biochemical experiments show that the Mut3 substitutions not only perturb the previously described Pat1-binding interface^46^, but also strongly impair RNA binding and Dhh1 condensation *in vitro*. These findings therefore provide additional biochemical context for the previously reported cellular Mut3 phenotype and indicate that its effects are not explained solely by disruption of the Dhh1–Pat1 interaction.

## DISCUSSION

This study establishes sequential Design of Experiments (DoE) as a framework for resolving how multiple biochemical variables jointly shape biomolecular condensation. The workflow progresses from screening, which prioritises influential variables, to factorial analysis, which quantifies interactions, and response-surface modelling, which resolves curvature and nonlinear behaviour within a selected experimental space. Each stage uses the information obtained previously to determine where additional experimental resolution is most valuable. The principal advantage is therefore not simply a reduction in the number of conditions, but the allocation of limited resources and measurements towards interpretable decisions, independent validation and mechanistically informative regions of a multidimensional landscape.

Sequential DoE occupies a complementary position between conventional one-factor-at-a-time (OFAT) experiments and dense high-throughput phase mapping. OFAT experiments are intuitive and effective for testing defined hypotheses, but they sample narrow trajectories through experimental space and cannot directly estimate interactions. Microfluidic and combinatorial platforms can provide substantially denser coverage, but may require significantly more protein and specialised experimental infrastructure^25,26,29^. Sequential DoE instead emphasises staged statistical prioritisation and can be implemented using standard biochemical reconstitution and fluorescence microscopy. The resulting models are not exhaustive phase diagrams or thermodynamic descriptions of phase boundaries, nor do they provide precise predictions for every condition. Instead, they provide representations of the principal response patterns within defined experimental ranges and identify relationships that warrant targeted validation.

Applying this workflow to Dhh1, Ded1 and PGL-3 revealed that the same biochemical inputs can generate distinct condensation landscapes. Ded1 provided a particularly clear example: the presence of poly(U) suppressed condensation at low NaCl but promoted or restored condensation as NaCl increased. An experiment performed at only one salt concentration could therefore have classified RNA as either inhibitory or promoting, depending on the background condition selected. Individual variables should consequently not be described as universally promoting or inhibiting condensation since their apparent effects can emerge from interactions with the surrounding biochemical environment.

The co-condensation experiments extended this principle to multicomponent assemblies. From the different interaction patterns of the proteins and their condensate-in-condensate morphologies, it became clear that influences on bulk condensation and changes in internal organisation are distinct properties of multicomponent systems and cannot necessarily be inferred from one another.

These contrasting behaviours suggest a possible relationship between condensate miscibility and cross-protein regulation. Our experiments showed that Dhh1 and Ded1 formed homogeneously mixed condensates when paired, whereas pairing Ded1 and Dhh1 with PGL-3 resulted in condensate-within-condensate assemblies. One possible explanation is that heterotypic interactions between Ded1 and Dhh1^51,52^, consistent with the pairwise interaction detected by the DoE analysis, favour mixing and allow one protein to influence the condensation response of the other. In contrast, preferentially homotypic interactions (i.e. no detectable influence of the secondary protein on the primary protein) may favour demixing and allow each of the proteins to retain a more independent response landscape. Differences in the molecular determinants of condensation might contribute to these behaviours: PGL-3 condensation is primarily driven by oligomerization of its folded D1–D2 domains^53^, whereas Dhh1 and Ded1 depend strongly on contributions from their disordered regions^14,16,43^. This interpretation remains speculative, as the DoE analysis does not directly measure molecular interaction strengths. Direct measurements of partitioning, interaction strengths and component dynamics, together with domain-specific perturbations, will be required to determine how miscibility relates to cross-protein regulation.

The Dhh1 findings illustrate how DoE can convert an unexpected response into a mechanistic hypothesis. ATP did not promote Dhh1 condensation within the investigated design space, despite the established coupling of ATP binding to canonical RNA engagement and conformational cycling in DEAD-box ATPases^16,37,43–45^. Instead, our follow-up experiments showed that Dhh1 can condense without added ATP and linked this behaviour to an ATP-independent RNA-binding surface on RecA2. One possible explanation for differences from previous studies is that ATP addition can alter assay pH if not rigorously controlled. This is particularly relevant given the strong pH dependence of Dhh1 condensation identified here. Under the conditions tested, the secondary RecA2 RNA-binding surface therefore appears to contribute more strongly to condensation than the canonical ATP-dependent RNA-binding mode, suggesting that the relative importance of the two RNA-binding modes may differ between the helicase cycle and condensate assembly.

The secondary RNA binding surface was previously also implicated in protein-protein interactions, e.g. with EDC3 and PAT1 to enhance PB formations^48^. Our data suggest that this is even more multifunctional than previously suggested, and that RNA binding and protein-mediated interactions act cooperatively to tune Dhh1 condensation and adapt PB formation to cellular conditions, e.g. high - low cellular RNA.

Several limitations constrain the conclusions that can be drawn from the here-established DoE analysis of condensate behaviour. The response surfaces are empirical models restricted to the proteins and conditions examined, and significant lack of fit for some models indicates that local variation remains unresolved. If relevant for the experimental systems, this should be addressed by targeted validation. Moreover, the coefficient of variation reports spatial heterogeneity in fluorescence intensity and is therefore a practical proxy for condensation rather than a direct measurement of phase boundaries or material properties. For systems in which more specialised properties are relevant, appropriate readouts should replace CV as the response variable in the DoE analysis. In sum, the framework should consequently be used to identify reproducible global relationships and prioritise follow-up experiments, not to replace orthogonal biophysical validation.

Together, these results show that, using standard equipment and relatively little sample, sequential DoE can extract substantial information from a limited number of experiments, and detect context-dependent condensation behaviour that would be difficult to recognise from isolated measurements alone. The ability to test many potential variables with little additional experimental effort will enable a more comprehensive understanding of biomolecular condensates that better accounts for their complex, dynamic nature.

## DATA AVAILABILITY

All data and analyses are included in main and supplementary figures. Raw microscopy data will be available in a public repository upon publication.

## CODE AVAILABILITY

Data was analysed using standard software, as described in methods and is available upon request.

## ACKNOWLEDGEMENTS

We thank all members of the Hondele laboratory for their thoughtful feedback on the manuscript and stimulating discussions throughout this work. We are also grateful to Debdatto Mookherjee for carefully reading the manuscript and providing valuable comments. We especially thank Timothy Sharp and Tobias Mühlethaler at the Biozentrum Biophysics Facility, as well as the staff of the Imaging Core Facility (IMCF) at the Biozentrum, University of Basel, for their expertise, advice and support.

## AUTHOR CONTRIBUTIONS

Conceptualization, M.S., and M.H.;

methodology, M.S.

formal analysis, M.S, and F.W.;

investigation, M.S, and F.W;

resources, M.S, C.N, and F.W;

Writing - original draft, M.S., F.W. and M.H.;

visualization, M.S, and F.W;

supervision, M.S, and M.H;

funding acquisition, M.H.

## FUNDING

M.H. acknowledges funding by the Swiss National Science Foundation [PCEFP3_187052] and the European Research Council [ERC Starting Grant 950262] and institutional support from the Kanton Basel-Stadt and Basel-Landschaft provided to the Biozentrum, University of Basel.

## COMPETING INTERESTS

The authors declare no competing interests.

## DECLARATION OF AI AND AI-A

During the preparation of this work, the author(s) used ChatGPT to improve wording and the clarity of the manuscript and to assist with script writing. The author(s) reviewed and edited the content as appropriate and take full responsibility for the content of the publication.

## Supplementary Information

**Supplementary Figure 1.**
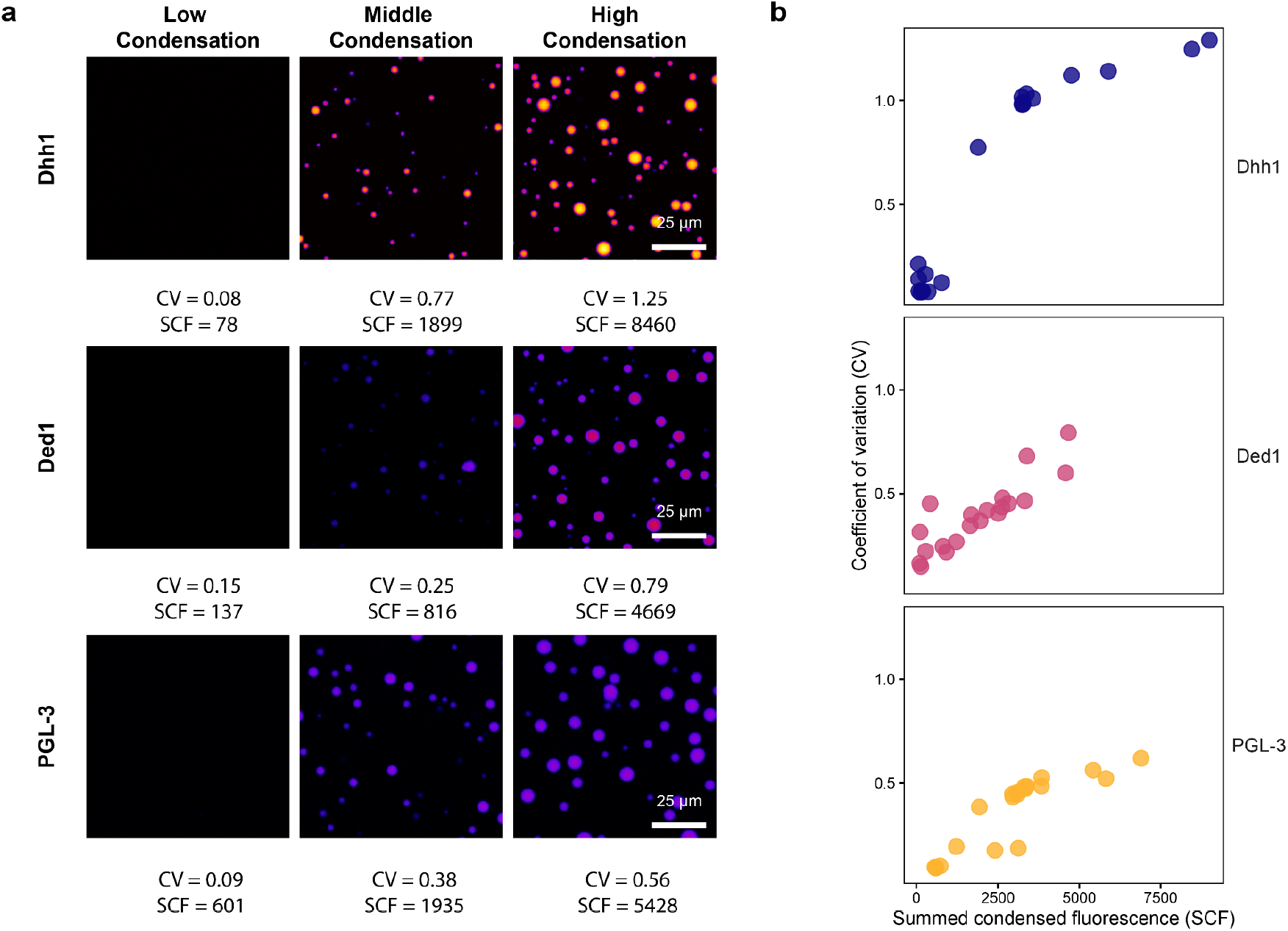
| Validation of the coefficient of variation as a proxy for condensation. Data were obtained from the central composite face-centred (CCF) design-of-experiments study performed with 5 µM protein. **a** Representative fluorescence images of Dhh1, Ded1, and PGL-3 under conditions producing low, intermediate, and high levels of condensation. The corresponding mean coefficient of variation (CV) and mean summed condensed fluorescence (SCF) are indicated below each image. For each field of view, SCF was calculated by summing the integrated fluorescence intensities of all segmented condensates. CV and SCF were subsequently averaged across the analysed fields of view within each well. Increasing condensation is accompanied by higher CV and SCF values. Scale bars, 25 µm. **b** Relationship between SCF and CV at the well level for Dhh1, Ded1, and PGL-3. Each point represents the mean value for one well. CV increases monotonically with SCF across the investigated condensation range for all three proteins, although the magnitude and shape of the relationship differ between proteins. This consistent positive relationship supports the use of CV as a high-throughput proxy for condensation-associated spatial heterogeneity.

**Supplementary Figure 2.**
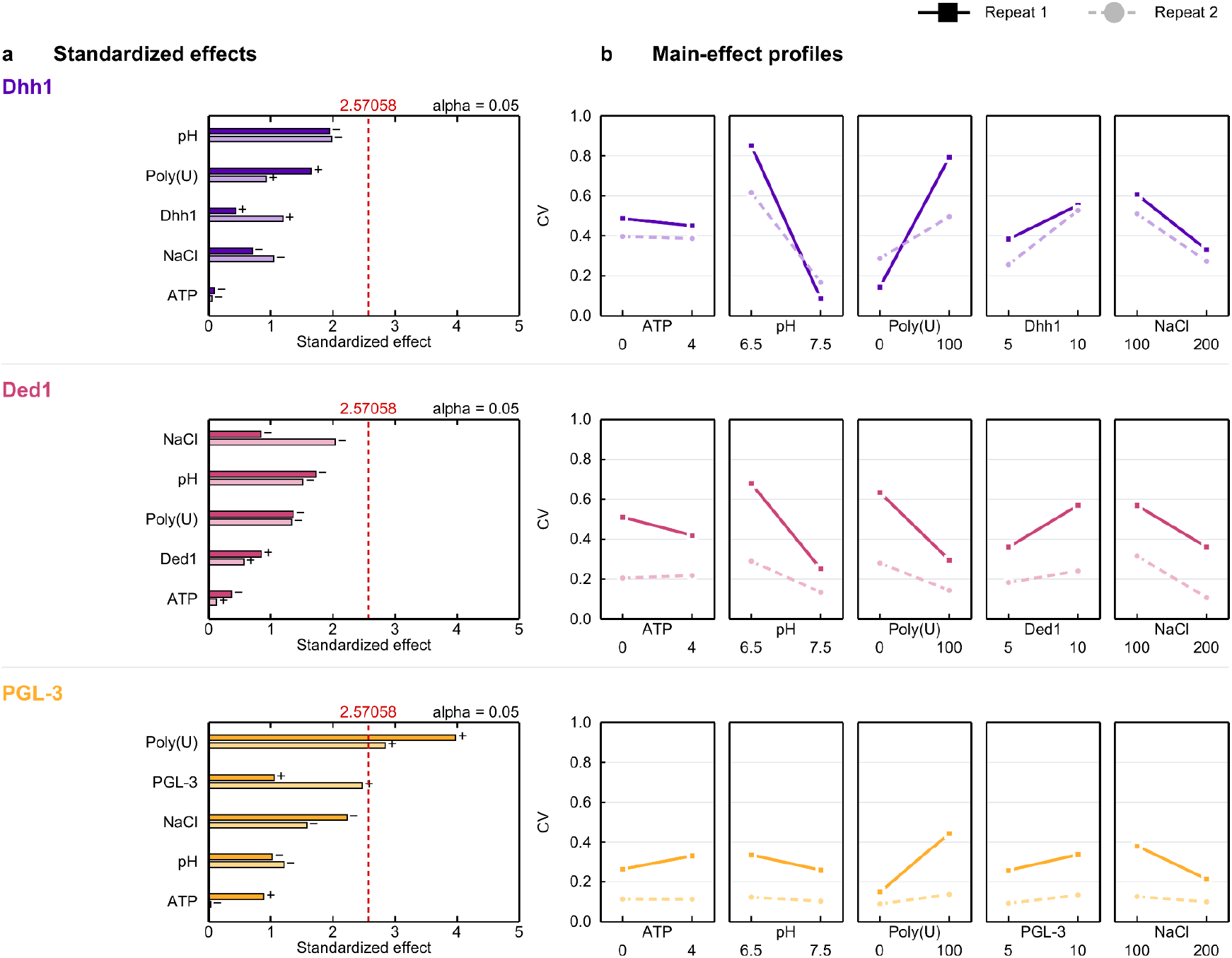
| Comparison of experimental repeats in the definitive screening design. **a** Repeat-specific Pareto plots showing the standardized effects of ATP, pH, Poly(U), protein concentration and NaCl on the coefficient of variation (CV) responses of Dhh1, Ded1 and PGL-3. Dark bars represent Repeat 1 and lighter bars represent Repeat 2; plus and minus signs indicate the direction of each effect. The red dashed line marks the critical standardized effect at α = 0.05. **b** Corresponding main-effect profiles across the investigated factor ranges: ATP, 0–4 mM; pH, 6.5–7.5; Poly(U), 0–100 µg/mL; protein, 5–10 µM; and NaCl, 100–200 mM. Repeat 1 is shown by solid lines and square markers and Repeat 2 by dashed lines and circular markers. All main-effect panels use the same CV scale of 0–1. Dhh1, Ded1 and PGL-3 are shown in purple, magenta and orange, respectively. The repeats showed different overall responses for PGL-3 (P < 0.001) and Ded1 (P = 0.015), but not for Dhh1 (P = 0.60). Because a different protein preparation was used in each repeat, the observed repeat effect cannot be attributed exclusively to protein preparation. Experimental repeat was therefore included as a blocking factor in the combined screening models.

**Supplementary Figure 3.**
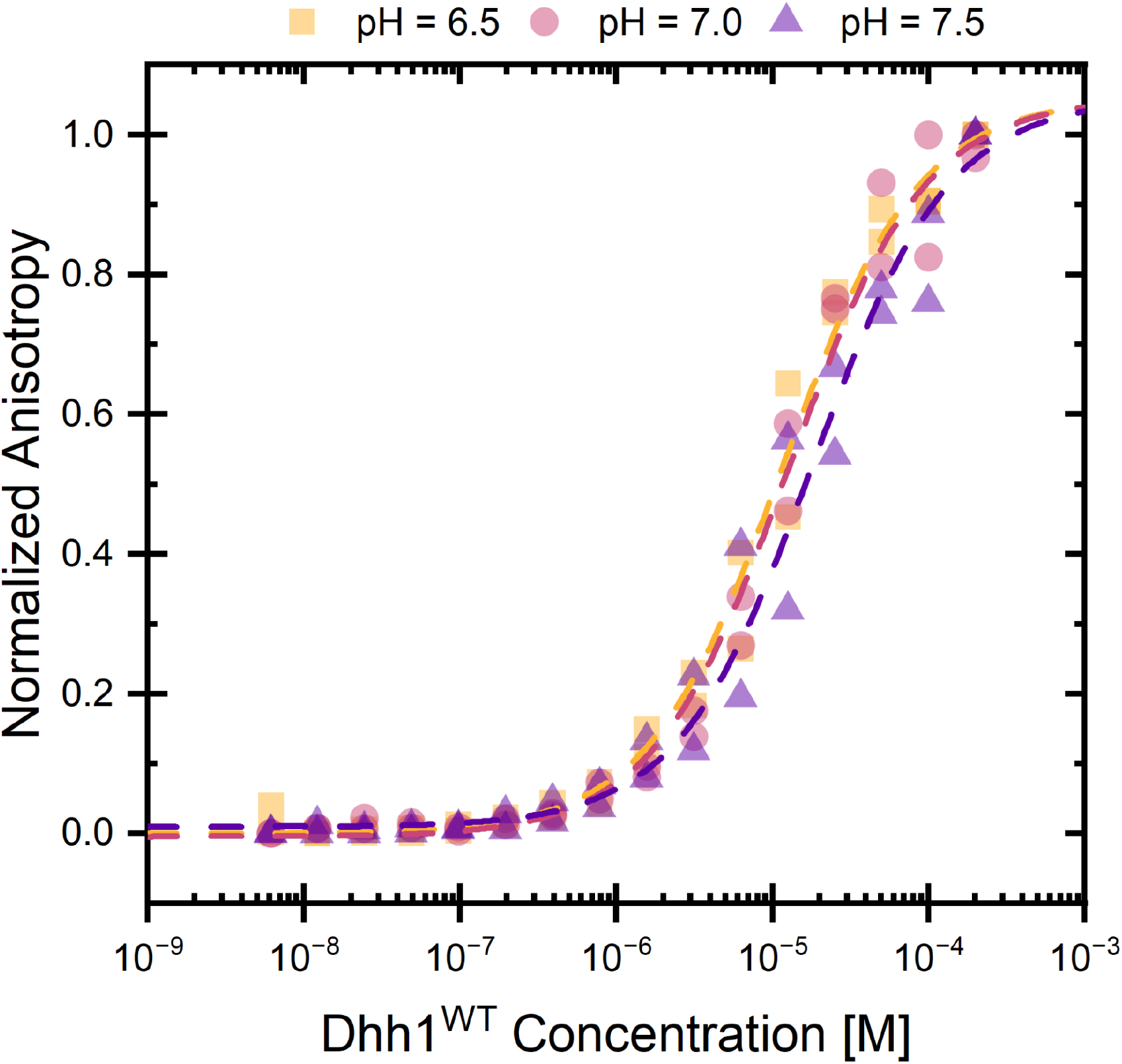
| Effect of pH on RNA binding by Dhh1^WT^. RNA binding was measured by fluorescence anisotropy as a function of Dhh1^WT^ concentration at pH 6.5 (yellow squares), pH 7.0 (red circles), and pH 7.5 (purple triangles), in the presence of 100 mM NaCl. Anisotropy values were normalized to the fitted minimum and maximum responses. Measurements were performed in duplicate (n = 2), with individual measurements shown. Dashed lines represent nonlinear fits to a binding isotherm. The binding curves largely overlapped, indicating that only minor differences in Dhh1^WT^ RNA binding were detected across the investigated pH range.

**Supplementary Figure 4.**
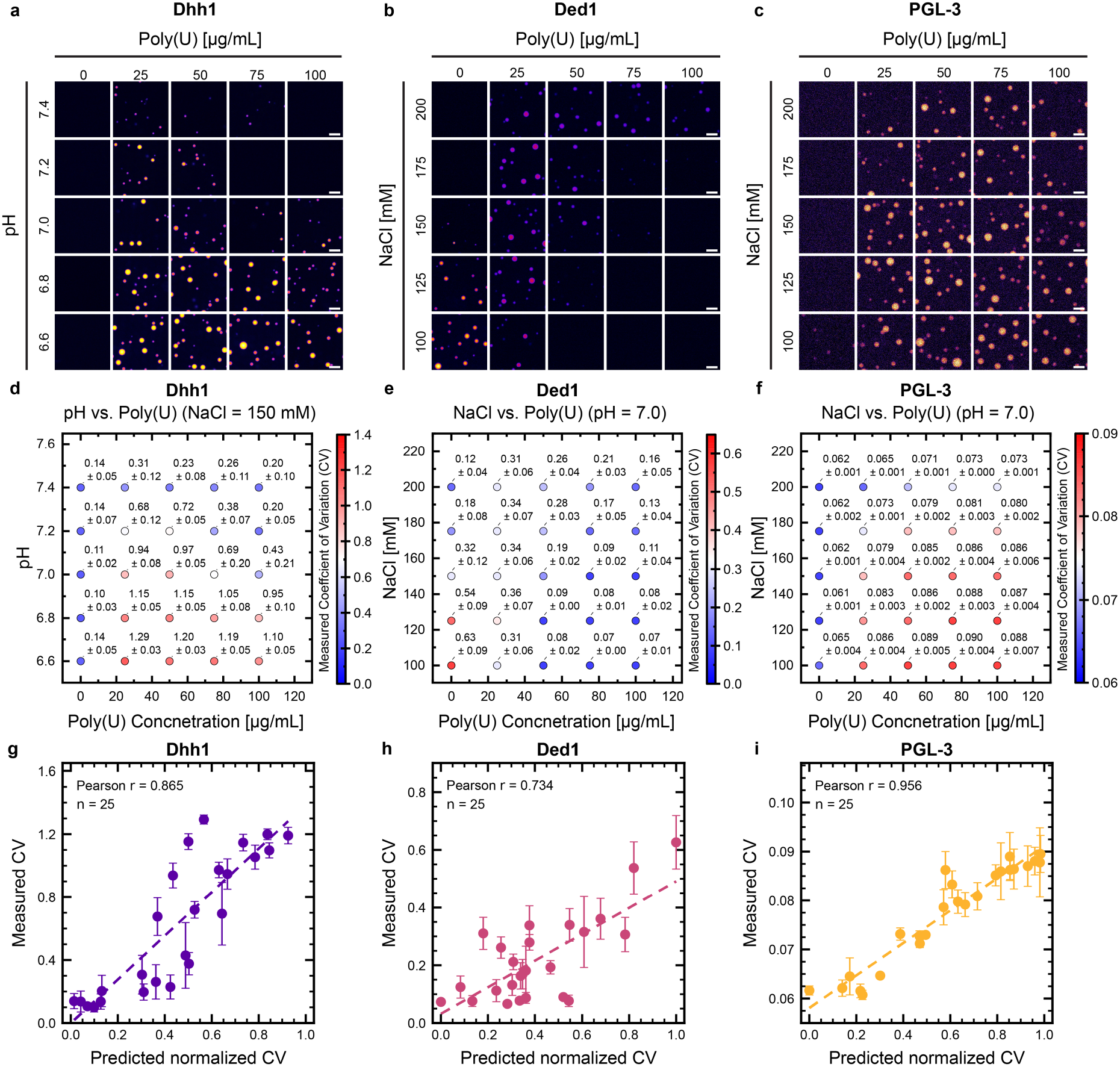
| Independent benchmarking of the CCF-derived condensation landscapes. **a** Representative fluorescence microscopy images of the Dhh1 benchmarking experiment. Conditions: 0–100 µg/mL poly(U), pH 6.6–7.4 and 150 mM NaCl. **b** Representative fluorescence microscopy images of the Ded1 benchmarking experiment. Conditions: 0–100 µg/mL poly(U), 100–200 mM NaCl and pH 7.0. **c** Representative fluorescence microscopy images of the PGL-3 benchmarking experiment. Conditions: 0–100 µg/mL poly(U), 100–200 mM NaCl and pH 7.0. Protein concentration was fixed at 5 µM in all experiments. Brightness and contrast settings were applied identically to all images within each panel but adjusted independently between proteins to accommodate differences in fluorescence intensity. Image adjustments were applied only for visualization; CV quantification was performed using the unadjusted images. **d** Quantification of the Dhh1 benchmarking experiment using the coefficient of variation (CV) of the protein fluorescence signal. **e** Quantification of the Ded1 benchmarking experiment. **f** Quantification of the PGL-3 benchmarking experiment. For **d-f** circle colour represents the mean measured CV and values above each condition show mean ± SD from three independently assembled reactions (n = 3). For each reaction the reaction-level CV was calculated as the mean CV across four fields of view. **g-i** Correlation between CCF model predictions and independent experimental measurements for Dhh1, Ded1 and PGL-3, respectively. Each point represents the mean measured CV ± SD from three independently assembled reactions, plotted against the predicted normalized CV at the corresponding condition. Each protein was evaluated across 25 validation conditions. Dashed lines show linear regression fits; Pearson’s correlation coefficients were r = 0.865 for Dhh1, r = 0.734 for Ded1 and r = 0.956 for PGL-3. Benchmarking measurements were not used to fit or select the models. Scale bars, 10 µm.

**Supplementary Figure 5.**
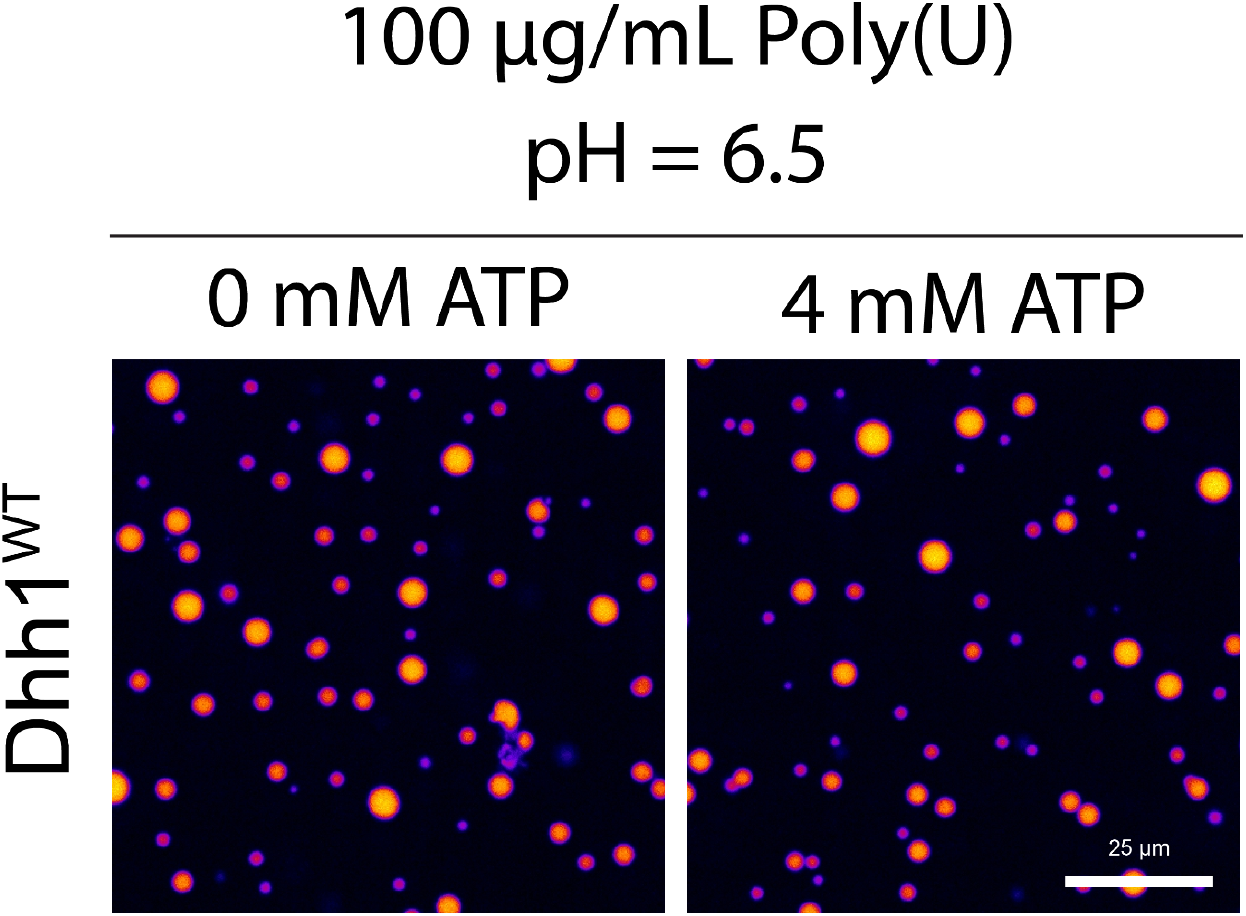
| ATP has minimal influence on Dhh1 condensation under the DoE conditions. Representative fluorescence microscopy images of Dhh1 condensation at 0 and 4 mM ATP in the presence of 100 µg/mL poly(U), 100 mM NaCl and at pH 6.5. Dhh1 formed abundant condensates at both ATP concentrations, showing that increasing ATP across the range used in the DoE screen has only a minor effect on condensation under these conditions. Scale bars, 25 µm.

**Supplementary Table 1.**
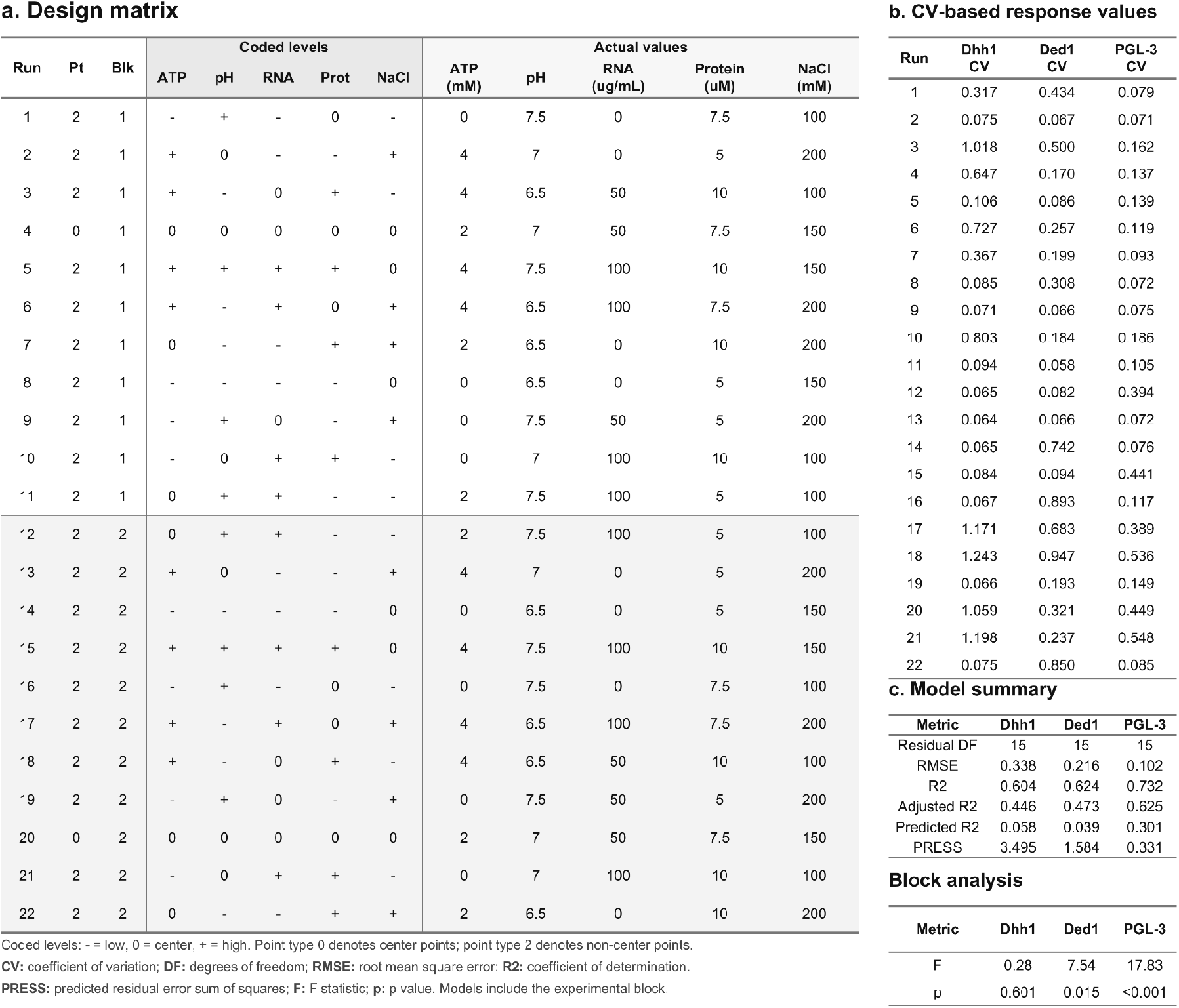
| Definitive screening design for Dhh1, Ded1 and PGL-3. **a** Design matrix for the 22-run definitive screening design used to evaluate the effects of ATP concentration, pH, poly(U) concentration, protein concentration and NaCl concentration. Coded factor levels and corresponding experimental values are shown separately. The experiment was performed in two blocks corresponding to two independent experimental repeats. **b** Coefficient of variation (CV) of the primary protein fluorescence channel for Dhh1, Ded1 and PGL-3 in each experimental condition. **c** Summary statistics for the main-effects models, including the experimental block as a model term. The block analysis reports the F statistic and corresponding p value for the difference between experimental blocks for each protein. CV, coefficient of variation; RMSE, root mean squared error; R2, coefficient of determination; PRESS, predicted residual error sum of squares.

a. Design matrix
| Run | Pt | Blk | Coded levels |  |  |  |  | Actual values |  |  |  |  |
| --- | --- | --- | --- | --- | --- | --- | --- | --- | --- | --- | --- | --- |
|  |  |  | ATP | pH | RNA | Prot | NaCl | ATP (mM) | pH | RNA (ug/mL) | Protein (uM) | NaCl (mM) |
| 1 | 2 | 1 | - | + | - | 0 | - | 0 | 7.5 | 0 | 7.5 | 100 |
| 2 | 2 | 1 | + | 0 | - | - | + | 4 | 7 | 0 | 5 | 200 |
| 3 | 2 | 1 | + | - | 0 | + | - | 4 | 6.5 | 50 | 10 | 100 |
| 4 | 0 | 1 | 0 | 0 | 0 | 0 | 0 | 2 | 7 | 50 | 7.5 | 150 |
| 5 | 2 | 1 | + | + | + | + | 0 | 4 | 7.5 | 100 | 10 | 150 |
| 6 | 2 | 1 | + | - | + | 0 | + | 4 | 6.5 | 100 | 7.5 | 200 |
| 7 | 2 | 1 | 0 | - | - | + | + | 2 | 6.5 | 0 | 10 | 200 |
| 8 | 2 | 1 | - | - | - | - | 0 | 0 | 6.5 | 0 | 5 | 150 |
| 9 | 2 | 1 | - | + | 0 | - | + | 0 | 7.5 | 50 | 5 | 200 |
| 10 | 2 | 1 | - | 0 | + | + | - | 0 | 7 | 100 | 10 | 100 |
| 11 | 2 | 1 | 0 | + | + | - | - | 2 | 7.5 | 100 | 5 | 100 |
| 12 | 2 | 2 | 0 | + | + | - | - | 2 | 7.5 | 100 | 5 | 100 |
| 13 | 2 | 2 | + | 0 | - | - | + | 4 | 7 | 0 | 5 | 200 |
| 14 | 2 | 2 | - | - | - | - | 0 | 0 | 6.5 | 0 | 5 | 150 |
| 15 | 2 | 2 | + | + | + | + | 0 | 4 | 7.5 | 100 | 10 | 150 |
| 16 | 2 | 2 | - | + | - | 0 | - | 0 | 7.5 | 0 | 7.5 | 100 |
| 17 | 2 | 2 | + | - | + | 0 | + | 4 | 6.5 | 100 | 7.5 | 200 |
| 18 | 2 | 2 | + | - | 0 | + | - | 4 | 6.5 | 50 | 10 | 100 |
| 19 | 2 | 2 | - | + | 0 | - | + | 0 | 7.5 | 50 | 5 | 200 |
| 20 | 0 | 2 | 0 | 0 | 0 | 0 | 0 | 2 | 7 | 50 | 7.5 | 150 |
| 21 | 2 | 2 | - | 0 | + | + | - | 0 | 7 | 100 | 10 | 100 |
| 22 | 2 | 2 | 0 | - | - | + | + | 2 | 6.5 | 0 | 10 | 200 |

b. CV-based response values
| Run | Dhh1 CV | Ded1 CV | PGL-3 CV |
| --- | --- | --- | --- |
| 1 | 0.317 | 0.434 | 0.079 |
| 2 | 0.075 | 0.067 | 0.071 |
| 3 | 1.018 | 0.500 | 0.162 |
| 4 | 0.647 | 0.170 | 0.137 |
| 5 | 0.106 | 0.086 | 0.139 |
| 6 | 0.727 | 0.257 | 0.119 |
| 7 | 0.367 | 0.199 | 0.093 |
| 8 | 0.085 | 0.308 | 0.072 |
| 9 | 0.071 | 0.066 | 0.075 |
| 10 | 0.803 | 0.184 | 0.186 |
| 11 | 0.094 | 0.058 | 0.105 |
| 12 | 0.065 | 0.082 | 0.394 |
| 13 | 0.064 | 0.066 | 0.072 |
| 14 | 0.065 | 0.742 | 0.076 |
| 15 | 0.084 | 0.094 | 0.441 |
| 16 | 0.067 | 0.893 | 0.117 |
| 17 | 1.171 | 0.683 | 0.389 |
| 18 | 1.243 | 0.947 | 0.536 |
| 19 | 0.066 | 0.193 | 0.149 |
| 20 | 1.059 | 0.321 | 0.449 |
| 21 | 1.198 | 0.237 | 0.548 |
| 22 | 0.075 | 0.850 | 0.085 |

c. Model summary
| Metric | Dhh1 | Ded1 | PGL-3 |
| --- | --- | --- | --- |
| Residual DF | 15 | 15 | 15 |
| RMSE | 0.338 | 0.216 | 0.102 |
| R2 | 0.604 | 0.624 | 0.732 |
| Adjusted R2 | 0.446 | 0.473 | 0.625 |
| Predicted R2 | 0.058 | 0.039 | 0.301 |
| PRESS | 3.495 | 1.584 | 0.331 |

| Metric | Dhh1 | Ded1 | PGL-3 |
| --- | --- | --- | --- |
| F | 0.28 | 7.54 | 17.83 |
| p | 0.601 | 0.015 | <0.001 |

**Supplementary Table 2.**
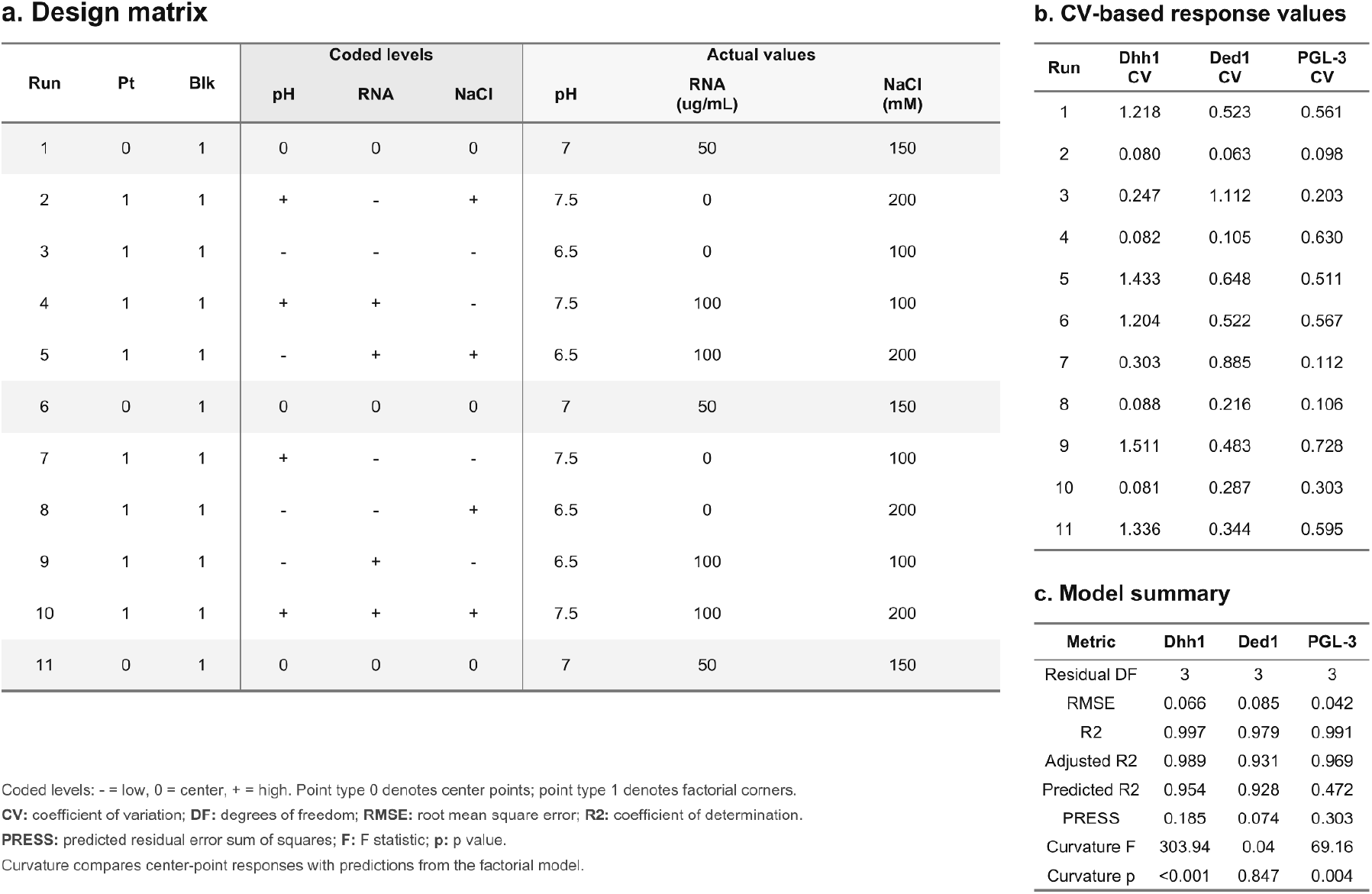
| Two-level full-factorial design for Dhh1, Ded1 and PGL-3. **a** Design matrix for the 11-run two-level full-factorial experiment used to investigate the effects of pH, poly(U) concentration and NaCl concentration. The design comprised eight factorial corner conditions and three centre-point replicates. Coded factor levels and corresponding experimental values are shown separately. **b** Coefficient of variation of the primary protein fluorescence channel for Dhh1, Ded1 and PGL-3 in each experimental condition. **c** Summary statistics for factorial models containing the three main effects and all two-factor interactions. RMSE, root-mean-square error; R², coefficient of determination; adjusted R², R² corrected for model complexity; predicted R², cross-validation-based estimate of predictive performance; PRESS, predicted residual error sum of squares.

**Supplementary Table 3.** | Central composite face-centred design for Dhh1, Ded1 and PGL-3. **a** Design matrix for the 20-run central composite face-centred experiment used to investigate the response surface defined by pH, poly(U) concentration and NaCl concentration. The design comprised eight cube points, six axial points and six centre-point replicates. Coded factor levels and corresponding experimental values are shown separately. **b** Coefficient of variation of the primary protein fluorescence channel for Dhh1, Ded1 and PGL-3 in each experimental condition. **c** Comparison of the full quadratic and reduced hierarchical response-surface models. Model summaries report residual degrees of freedom, RMSE, R², adjusted R², predicted R², PRESS and the lack-of-fit *p* value for each protein.

a. Design matrix
| Run | Pt | Blk | Coded levels |  |  | Actual values |  |  |
| --- | --- | --- | --- | --- | --- | --- | --- | --- |
|  |  |  | NaCl | pH | RNA | NaCl (mM) | pH | RNA (ug/mL) |
| 1 | -1 | 1 | 0 | - | 0 | 150 | 6.5 | 50 |
| 2 | 0 | 1 | 0 | 0 | 0 | 150 | 7 | 50 |
| 3 | -1 | 1 | + | 0 | 0 | 200 | 7 | 50 |
| 4 | 0 | 1 | 0 | 0 | 0 | 150 | 7 | 50 |
| 5 | -1 | 1 | 0 | 0 | + | 150 | 7 | 100 |
| 6 | 1 | 1 | + | - | + | 200 | 6.5 | 100 |
| 7 | 1 | 1 | + | + | + | 200 | 7.5 | 100 |
| 8 | 1 | 1 | - | + | - | 100 | 7.5 | 0 |
| 9 | 1 | 1 | + | + | - | 200 | 7.5 | 0 |
| 10 | 1 | 1 | - | - | + | 100 | 6.5 | 100 |
| 11 | 1 | 1 | - | - | - | 100 | 6.5 | 0 |
| 12 | 1 | 1 | + | - | - | 200 | 6.5 | 0 |
| 13 | -1 | 1 | 0 | + | 0 | 150 | 7.5 | 50 |
| 14 | -1 | 1 | 0 | 0 | - | 150 | 7 | 0 |
| 15 | 0 | 1 | 0 | 0 | 0 | 150 | 7 | 50 |
| 16 | 0 | 1 | 0 | 0 | 0 | 150 | 7 | 50 |
| 17 | 0 | 1 | 0 | 0 | 0 | 150 | 7 | 50 |
| 18 | -1 | 1 | - | 0 | 0 | 100 | 7 | 50 |
| 19 | 0 | 1 | 0 | 0 | 0 | 150 | 7 | 50 |
| 20 | 1 | 1 | - | + | + | 100 | 7.5 | 100 |

b. CV-based response values
| Run | Dhh1 CV | Ded1 CV | PGL-3 CV |
| --- | --- | --- | --- |
| 1 | 1.291 | 0.600 | 0.526 |
| 2 | 1.033 | 0.370 | 0.447 |
| 3 | 0.080 | 0.406 | 0.176 |
| 4 | 1.010 | 0.346 | 0.455 |
| 5 | 0.775 | 0.220 | 0.485 |
| 6 | 1.122 | 0.466 | 0.385 |
| 7 | 0.075 | 0.269 | 0.194 |
| 8 | 0.082 | 0.681 | 0.101 |
| 9 | 0.082 | 0.150 | 0.091 |
| 10 | 1.247 | 0.399 | 0.619 |
| 11 | 0.214 | 0.794 | 0.186 |
| 12 | 0.088 | 0.316 | 0.096 |
| 13 | 0.124 | 0.224 | 0.432 |
| 14 | 0.141 | 0.453 | 0.094 |
| 15 | 0.983 | 0.421 | 0.484 |
| 16 | 0.992 | 0.479 | 0.484 |
| 17 | 1.016 | 0.453 | 0.446 |
| 18 | 1.142 | 0.246 | 0.563 |
| 19 | 0.983 | 0.436 | 0.475 |
| 20 | 0.164 | 0.165 | 0.522 |

c. Model summary Full model
| Metric | Dhh1 | Ded1 | PGL-3 |
| --- | --- | --- | --- |
| DF | 10 | 10 | 10 |
| RMSE | 0.264 | 0.099 | 0.054 |
| R<sup>2</sup> | 0.847 | 0.813 | 0.953 |
| Adjusted R<sup>2</sup> | 0.710 | 0.644 | 0.910 |
| Predicted R<sup>2</sup> | -0.296 | -0.236 | 0.501 |
| PRESS | 4.825 | 0.652 | 0.271 |
| LOF p | <0.001 | 0.029 | 0.003 |

| Metric | Dhh1 | Ded1 | PGL-3 |
| --- | --- | --- | --- |
| DF | 14 | 15 | 13 |
| RMSE | 0.245 | 0.086 | 0.054 |
| R<sup>2</sup> | 0.816 | 0.790 | 0.938 |
| Adjusted R<sup>2</sup> | 0.751 | 0.734 | 0.910 |
| Predicted R<sup>2</sup> | 0.651 | 0.636 | 0.759 |
| PRESS | 1.597 | 0.191 | 0.129 |
| LOF p | <0.001 | 0.077 | 0.004 |

**Supplementary Table 4.**
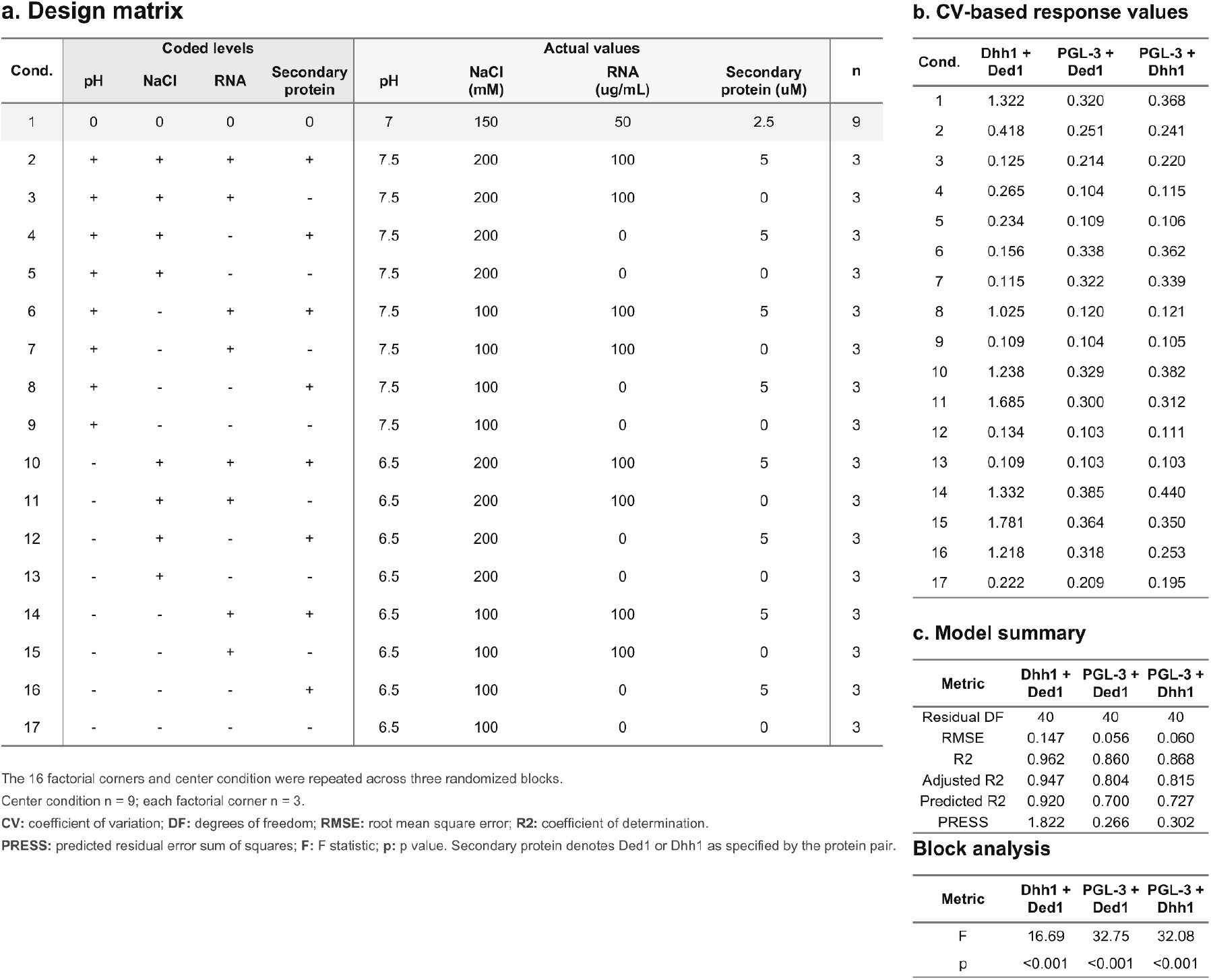
| Factorial analysis of the co-condensation assays. **a** Shared design matrix used for the Dhh1–Ded1, Ded1–PGL-3 and Dhh1–PGL-3 co-condensation experiments. The effects of pH, NaCl concentration, poly(U) concentration and client-protein concentration were investigated using 16 factorial corner conditions and one centre condition across three randomized experimental blocks. Coded factor levels and corresponding experimental values are shown separately. The experiment comprised three independent blocks. Each factorial corner was measured once per block, whereas the centre condition was measured three times within each block. **b** Coefficient of variation of the primary protein fluorescence channel for each protein pair, averaged across the three independent experimental blocks. **c** Model and block summary reporting residual degrees of freedom, RMSE, R², adjusted R², predicted R², PRESS and the omnibus block *F* and *p* values. Significant differences between blocks were detected for all three protein pairs, with *p* < 0.001. Client denotes Ded1 or Dhh1, as specified by the protein pair. RMSE, root-mean-square error; R², coefficient of determination; predicted R², cross-validation-based estimate of predictive performance; PRESS, predicted residual error sum of squares.

**Supplementary Table 5.** | Key resources.

| REAGENT, RESOURCE or TOOL | SOURCE | IDENTIFIER |
| --- | --- | --- |
| **Bacterial Strains** |  |  |
| E. coli Lemo21(DE3) | NEB | Cat# C2528 |
| **Antibodies** |  |  |
| Mouse anti-V5 | GeneTex | Cat# GTX635260 |
| Mouse anti-Pgk1 | Abcam | Cat# ab113687 |
| HRP-conjugated anti-mouse IgG secondary antibody | Sigma-Aldrich | Cat# A0168 |
| **Chemicals and Solutions** |  |  |
| 2-ME; 2-mercaptoethanol | Carl Roth | Cat# 4227.3 |
| Ampicillin sodium salt | Carl Roth | Cat# K029.2 |
| ATP; Adenosine 5'-triphosphate disodium salt hydrate | Merck | Cat# A2383 |
| ATTO 488 NHS-ester | ATTO-TEC | Cat# AD 488-31 |
| ATTO 565 NHS-ester | ATTO-TEC | Cat# AD 565-35 |
| BSA; Bovine serum albumin | Merck | Cat# A9418 |
| Chloramphenicol | Carl Roth | Cat# 3886.1 |
| Cyanine-5 hydrazide | Lumiprobe | Cat# 23070 |
| DNaseI | Applichem | Cat# A3778 |
| D(+)-glucose | Carl Roth | Cat# HN06.3 |
| Di-sodium hydrogen phosphate | Applichem | Cat# A1046 |
| D(+)-sucrose | Carl Roth | Cat# 9286.1 |
| DTT; Dithiothreitol | Applichem | Cat# A1101,0025 |
| Ethanol absolute | Merck | Cat# 1.00983 |
| Glycerol | Fisher<br>Bioreagents | Cat# BP229-4 |
| Imidazole | Carl Roth | Cat# 3899.4 |
| Immobilon Western chemiluminescent HRP<br>substrate | Merck | Cat# WBKLS0500 |
| Iodoacetamide | Applichem | Cat# A1666 |
| IPTG; Isopropyl-β-D-1-thiogalactopyranosid | LuBio | Cat# LU5002-0025 |
| Kanamycin sulphate | Carl Roth | Cat# T832.3 |
| Magnesium chloride hexahydrate | Carl Roth | Cat# 2189.1 |
| NucleoSpin Plasmid, Mini Prep Kit | Macherey Nagel | Cat# 740588.50 |
| PreScission Protease; C3-Protease | GeneScript | Cat# Z02799 |
| Prestained protein marker | Proteintech | Cat# PL00001 |
| Poly(U); Polyuridylic acid potassium salt | Merck | Cat# P9528 |
| Q5® High-Fidelity PCR Kit | NEB | Cat# E0555L |
| RedSafe™ Nucleic Acid Stain | iNtRON Bio | Cat# 21141 |
| RNase A | Macherey Nagel | Cat# 740505 |
| RNasin® Ribonuclease Inhibitor, recombinant | Promega | Cat# N2515 |
| Sodium acetate | Merck | Cat# 1062682500 |
| Sodium chloride | Carl Roth | Cat# 3957.1 |
| SDS; Sodium dodecyl sulphate | Carl Roth | Cat# CN30.3 |
| Sodium dihydrogen phosphate monohydrate | Merck | Cat# 1.06346 |
| Sodium metaperiodate | Merck | Cat# 1065970050 |
| T4 Ligase | in house | N/A |
| T4 PNK | NEB | Cat# M0201L |
| Tris; Tris(hydroxymethyl)aminomethane | Merck | Cat# T1503 |
| **Plasmid DNA** |  |  |
| pMH1539_pETMCN_His10-MPB_3C_Dhh1-WT | This Study | pMH1539 |
| pMH1540_pETMCN_His10-MPB_3C_Ded1_WT | This Study | pMH1540 |
| pMH1541_pETMCN_KAN_His10MBP_3C_PGL3 | This Study | pMH1541 |
| pMH1672_pETMCN_His10-MPB_3C_Dhh1_N269A_H272A_T276A_K280A (Mut1) | This Study | pMH1672 |
| pMH1673_pETMCN_His10-MPB_3C_Dhh1_S292D_N294D_R295D (Mut3) | This Study | pMH1673 |
| pMH1674_pETMCN_His10-MPB_3C_Dhh1[252-506] | This Study | pMH01674 |
| pMH1931_pETMCN_His10-MPB_3C_Dhh1[252-506] _Mut3 | This Study | pMH01931 |
| pMH2244_pETMCN_KAN_His10-TS_3C_Pat1[5-49] | Sachdev,<br>Ruchika, et al.<br>"Pat1 promotes processing body assembly by enhancing the phase separation of the DEAD-box ATPase Dhh1 and RNA." *elife* 8 (2019): e41415. | pMH2244 (pKW4530) |
| **Other Material** |  |  |
| 384-well, black, clear flat bottom plate | Corning | Cat# 3764 |
| 384-well, black, optically clear flat bottom, ultra-low-attachment-coated PhenoPlate | Revvity | Cat# 6057800 |
| MicroAmp Fast Optical 96-well plate | Applied Biosystemy | Cat# 4346907 |
| Amicon® Ultra Centrifugal Filter, 100 kDa MWCO | Merck | Cat# UFC910008 |
| Amicon® Ultra Centrifugal Filter, 30 kDa MWCO | Merck | Cat# UFC903008 |
| Amicon® Ultra Centrifugal Filter, 10 kDa MWCO | Merck | Cat# UFC901008 |
| Amicon® Ultra Centrifugal Filter, 3 kDa MWCO | Merck | Cat# UFC500396 |
| HiLoad® 26/600 Superdex® 200 pg | Cytiva | Cat# 28-9893-36 |
| **Software and Algorithms** |  |  |
| Adobe Illustrator V29.7.1 | Adobe | https://www.adobe.com/products/illustrator.html |
| CellProfiler™ V4.2.6 | (Stirling 2021) | https://cellprofiler.org |
| ChatGPT | OpenAI | https://chatgpt.com/ |
| OriginPro 2026 | OriginLab | https://www.originlab.com/2026 |
| NIS-Elements Software V5.42.07 | Nikon | N/A |
| OMERO server and clients | (Allan 2012) | https://www.openmicroscopy.org/omero/ |
| R Studio V4.5.3 | Posit team 2025 | https://posit.co/ |
| SnapGene software V8.2 | SnapGene | https://www.snapgene.com/ |
| Unicorn Software V7.13 | Cytiva | N/A |
| UniProt data base |  | www.uniprot.org |
| **Machines and Microscopes** |  |  |
| ÄKTA pure | GE Healthcare | N/A |
| EmulsiFlex | Avestin | N/A |
| Fusion FX imaging system | Vilber | N/A |
| NanoDrop Spectrophotometer | Thermo Scientific | N/A |
| Nikon Ti2 Crest Spinning Disk | Nikon | N/A |
| Odyssey M | LiCor | N/A |
| Spark Multi-Mode Plate Reader | Tecan | N/A |
| Ultracentrifuge | Beckmann Coulter | N/A |

**Supplementary Table 6.** | Yeast strains used in this study.

| Name | genotype | Source |
| --- | --- | --- |
| W303 | *MAT A ade2-1 can1-100 GAL phi+ his3-11,15 ura3-1 leu2-3, 112 trp1-1* |  |
| YMH2 | W303<br>*dhh1::DHH1-HIS3MX6 GFP dcp2::DCP2-mCherry-NatMX6* | KWY1973<br>Sachdev, R., Hondele, M., Linsenmeier, M., Vallotton, P., Mugler, C. F., Arosio, P., & Weis, K. (2019). Pat1 promotes processing body assembly by enhancing the phase separation of the DEAD-box ATPase Dhh1 and RNA. *eLife*, 8, e41415.<br>https://doi.org/10.7554/eLife.41415<br>Karsten Weis Lab |
| YMH565 | W303<br>*dhh1::KanMX6 dcp2::DCP2-mCherry-NatMX6* | KWY2193<br>Carroll, J. S., Munchel, S. E., & Weis, K. (2011). The DExD/H box ATPase Dhh1 functions in translational repression, mRNA decay, and processing body dynamics. *The Journal of cell biology*, 194(4), 527–537.<br>https://doi.org/10.1083/jcb.201007151 |
| YMH629<br>"Dhh1<sup>WT</sup>–V5" | W303<br>*dhh1Δ::KANMX dcp2::DCP2-mCherry-NAT*<br>*his3::4xlexA-pCYC1-DHH1-V5-HIS3MX6* | This study |
| YMH630<br>"Dhh1<sup>Mut3</sup>–V5" | W303<br>*dhh1Δ::KANMX dcp2::DCP2-mCherry-NAT*<br>*his3::4xlexA-pCYC1-DHH1-S292D,N294D,R295D-V5-HIS3MX6* | This study |
| YMH631<br>"Dhh1<sup>252-506</sup>–V5" | W303<br>*dhh1Δ::KANMX dcp2::DCP2-mCherry-NAT*<br>*his3::4xlexA-pCYC1-DHH1Δ4-753-V5-HIS3MX6* | This study |
| YMH632<br>"Dhh1<sup>252-506 + Mut3</sup>–V5" | W303<br>*dhh1Δ::KANMX dcp2::DCP2-mCherry-NAT*<br>*his3::4xlexA-pCYC1-DHH1Δ4-753-S292D,N294D,R295D-V5-HIS3MX6* | This study |

## METHODS

Reagents and resources are listed in Supplementary Table 5. Yeast strains are listed in Supplementary Table 6.

### Protein expression and purification

Recombinant proteins were expressed in *E. coli* Lemo21(DE3) from pETMCN-based expression plasmids. Overnight pre-cultures were grown in LB medium containing the appropriate antibiotics and 1% D-glucose. Expression cultures were inoculated in Terrific Broth medium supplemented with the appropriate antibiotics, grown at 37 °C to an OD600 of 0.6-0.8, and induced with 300 μM IPTG. After overnight expression at 18 °C, cells were harvested by centrifugation.

All subsequent purification steps were performed at 4 °C. Cell pellets were resuspended in lysis buffer (50 mM Tris-HCl pH 8.0, 1 M NaCl, 25 mM imidazole, 10% glycerol, protease inhibitors, DNase I and RNase A). Cells were lysed by pressure homogenisation followed by sonication, and lysates were clarified by centrifugation and filtration. His-tagged proteins were captured by immobilized metal affinity chromatography (IMAC) and eluted using an imidazole gradient up to 500 mM imidazole. Eluted proteins were subjected to 3C protease cleavage during overnight dialysis against dialysis buffer (50 mM Tris-HCl pH 7.5, 1 M NaCl, 25 mM imidazole, 2 mM MgCl2, 3 mM beta-mercaptoethanol, 10% glycerol). Cleaved affinity tags and uncleaved protein were removed by reverse IMAC. Proteins were further purified by size-exclusion chromatography using a HiLoad Superdex 200 pg 26/600 column equilibrated in SEC buffer (50 mM phosphate buffer pH 7.5, 1 M NaCl, 2 mM MgCl_2_, 3 mM beta-mercaptoethanol, 10% glycerol) at a flow rate of 2.6 mL/min. Clean SEC fractions were pooled, concentrated, aliquoted, snap-frozen in liquid nitrogen and stored at -80 °C. Protein purity was assessed by SDS-PAGE and Coomassie staining.

Proteins used for RNA-binding assays were additionally buffer-exchanged using a HiPrep 26/10 desalting column (Cytiva) into RNA-binding assay buffer (50 mM phosphate buffer pH 7.5, 200 mM NaCl, 2 mM MgCl_2_, 3 mM beta-mercaptoethanol, 10% glycerol, 10% 1,6-hexanediol).

### In vitro condensation assay

*In vitro* condensation assays were performed in 384-well black, optically clear, flat-bottom, ultra-low-attachment PhenoPlates (Revvity). Protein stocks were diluted in the corresponding protein storage buffer to ten times the final assay concentration. For fluorescence imaging, chemically labelled protein was mixed with unlabelled protein so that labelled protein constituted 1% of the final total protein concentration.

Condensation reactions were assembled by mixing 2 μL of tenfold-concentrated protein solution with 18 μL of the corresponding assay buffer, yielding a final reaction volume of 20 μL. Unless otherwise specified, reactions contained 2 mM MgCl₂, 0.5 mg/mL BSA and 1 mM DTT at final assay concentrations. The pH, NaCl concentration, RNA concentration and additional factors were adjusted according to the respective DoE condition or titration experiment. Poly(U) RNA was obtained from Sigma-Aldrich (P9528). Reactions were incubated for 30 min at 25 °C before imaging. Reaction assembly and image acquisition were staggered to maintain the same incubation time across all conditions within an experiment.

Images were acquired on a temperature-controlled inverted Nikon Ti2 microscope equipped with a Crest X-Light V3 spinning-disk confocal system, Nikon Perfect Focus System and Lumencor Celesta light source. Excitation/emission settings were 470 nm excitation with a 515/30 nm emission filter, 545 nm excitation with a 595/31 nm emission filter, and 640 nm excitation with a 698/70 nm emission filter. The microscope was controlled using Nikon NIS-Elements software, and image acquisition was automated using the integrated JOBS module. Four fields of view were acquired per well in the appropriate fluorescence channels. Exposure times and illumination intensities were kept constant within each experiment. Representative images were processed using linear brightness and contrast adjustments in OMERO; quantitative analysis was performed on raw images.

Images were analysed using CellProfiler (v4.2.6). For each field of view, condensate formation was quantified using the coefficient of variation (CV), calculated as the standard deviation of pixel intensities divided by the mean pixel intensity across the entire protein fluorescence image. CV values from all fields of view within the same well were averaged to obtain a mean CV for each well. For titration experiments, dose–response models were fitted in OriginPro 2026 (OriginLab) using the mean CV value from each independent biological replicate at each concentration as an individual observation.

### Design-of-experiments generation and analysis

DoE designs and statistical analyses were performed in OriginPro 2026 (OriginLab) using the Design of Experiments App (v2.79). Design matrices and factor levels are provided in the Supplementary Tables.

Each reaction was completely independently (no master mixes) assembled as a total volume of 20 µL in one well of a 384 well surface-coated plate (Revvity). Four fluorescence images were acquired from separate fields of view from each reaction well. The coefficient of variation (CV; standard deviation divided by the mean) of pixel intensities was calculated for each image, and the four image-level CVs were averaged to obtain a single reaction-level response.

Definitive screening designs (DSD) for Dhh1, Ded1 and PGL-3 contained 11 conditions and five factors: ATP, NaCl, pH, poly(U) and protein concentration. Each DSD was performed in two experimental blocks using independently purified and labelled protein preparations (22 observations per protein). Linear models contained the five coded main effects and experimental block; interaction and quadratic terms were not included. Because protein preparation differed between blocks, protein-preparation effects were confounded with block-associated variation. The blocks were additionally analysed separately to assess consistency of the main effects. Effects were standardized by their standard errors and visualized using Pareto plots. Statistical significance was evaluated at α = 0.05.

Following screening, pH, NaCl and poly(U) were analysed in a single experimental block using three-factor, two-level full factorial designs containing the eight possible combinations of low and high factor levels together with the centre condition prepared independently in triplicate (11 reactions in total). The corresponding models contained all three main effects, pairwise interactions and a centre-point indicator.

Face-centred central composite designs were subsequently performed in a single experimental block and contained 14 non-centre design conditions together with six replicated centre points (20 reactions in total). The models contained linear, interaction and quadratic terms for pH, NaCl and poly(U). Non-significant higher-order terms were removed while retaining all main effects and maintaining model hierarchy.

Model fit and predictive performance were assessed using R², adjusted R² and predicted R². R² describes the proportion of observed variation explained by the fitted model, whereas adjusted R² accounts for model complexity by penalizing the inclusion of additional terms that do not sufficiently improve model fit. Predicted R² estimates the model’s ability to predict observations not used during model fitting. Lack-of-fit tests and residual diagnostics were additionally used to assess model adequacy. For the central composite designs, the main response patterns inferred from the fitted models were further evaluated in independent benchmarking experiments performed at selected conditions within the investigated design space.

Co-condensation experiments were analysed using two-level factorial designs containing pH, NaCl, poly(U) and the secondary protein. Each experiment was performed in three separate blocks, and a centre-point indicator was included. PGL-3 CV was used as the response for PGL-3 scaffold mixtures and Dhh1 CV for Dhh1–Ded1 mixtures.

### RNA-binding fluorescence anisotropy assay

RNA binding was measured by fluorescence anisotropy using a 6-FAM-labelled 18-mer oligo(U) RNA. RNA master mix was prepared in assay buffer containing 50 mM phosphate buffer pH 7.0, 2 mM MgCl_2_, 2 mM DTT and 50 nM oligo(U)18-6-FAM. For each condition, 5 μL RNA master mix was transferred into a 384-well low-binding microplate (Corning, 3820) and mixed with 5 μL of the corresponding protein dilution by pipetting up and down five times. Protein dilution series was prepared in protein storage buffer specific for RNA binding.

Final assay conditions in the well were 25 nM oligo(U)18-6-FAM, 50 mM phosphate buffer pH 7.0, 100 mM NaCl, 2 mM MgCl2, 5% glycerol, 5% 1,6-hexanediol and 1 mM DTT. Reactions were incubated for 30 min at 25 °C before fluorescence anisotropy was measured using a Tecan Spark plate reader. Binding curves were generated by plotting fluorescence anisotropy as a function of protein concentration. n represents the number of independent replicates. All triplicate measurements are shown, and apparent dissociation constants were determined in OriginPro 2026 (OriginLab) by fitting all data points simultaneously to a one-site binding model.

### Chemical labelling of proteins and poly(U)

Purified recombinant proteins were chemically labelled with NHS ester dyes. PGL3 was labelled with ATTO488 NHS ester (ATTO-TEC), whereas Dhh1 and Ded1 were labelled with ATTO565 NHS ester (ATTO-TEC) for the single protein experiments. For the co-condensation experiments Ded1 was labelled with ATTO488 NHS ester in addition. To favour labelling of the N-terminal primary amine over lysine side chains, labelling reactions were performed at pH 6.5 or lower. PGL3 and ATTO488 NHS ester were mixed at a 1:4 molar ratio and incubated for 1 h at room temperature protected from light. Dhh1 or Ded1 and ATTO565 NHS ester were mixed at a 1:4 molar ratio and incubated overnight protected from light.

Free dye was removed by buffer exchange using HiTrap™ Desalting columns (Cytiva) into the corresponding storage buffer. Labelled proteins were concentrated using Amicon^®^ Ultra Centrifugal Filter (10 kDa MWCO, Merck), aliquoted, snap-frozen in liquid nitrogen and stored at -80 °C. Protein concentration and labelling efficiency were determined for each preparation. For microscopy-based LLPS assays, fluorescently labelled protein was spiked into the corresponding unlabelled protein at 1% of the final total protein concentration, unless stated otherwise.

Poly(U) was labelled at its 3′ end with Cyanine-5 hydrazide (Lumiprobe) using a modified protocol^54^. Reactions were prepared using DEPC-treated water and RNase-free tubes. Briefly, 22.3 µL of 10 mg/mL poly(U) was mixed with 10 µL of 0.25 M sodium acetate (Merck), pH 5.6, and water to a volume of 37.5 µL. Subsequently, 2.5 µL of 1 mM sodium metaperiodate (Merck) was added, and the reaction was incubated at 25 °C for 90 min in the dark. A further 2 µL of 2.5 mM sodium periodate was added, followed by incubation at 25 °C for 15 min. Finally, 3 µL of 2.5 mM Cyanine-5 hydrazide was added, and the reaction was incubated at 37 °C for 3 h in the dark. Labelled RNA was recovered by precipitation.

### Yeast imaging and P body quantification

Yeast cells were grown overnight in YPD at 30°C. Cultures were diluted to an OD_600_ of 0.2 in 750µl in a 2 ml Eppendorf tube and grown for 1h at 30°C and 1000 rpm in a tabletop Thermomixer. After 1h, either 750 µl of YPD or YPD with 1µM estradiol were added and the culture was grown for another 4h at 30°C and 1000 rpm. After 4h, the OD600 was measured and the amount of 1ml of OD600 of 0.1 was harvested by centrifugation (1 min at 8000 rcf). Pellets were resuspended in 500 µl of SC media with or without glucose, mixed by vortexing and pelleted again. The pellet was resuspended in 1 ml of SC media with or without glucose and 20 µl were pipetted into a 384-well plate and the remaining culture was used for TCA extraction and western blotting. Cells were pelleted in the plate for 30 s at 50 rcf.

Imaging of the Dcp2-mCherry signal was performed on an inverted Nikon Ti2 widefield fluorescence microscope using a 100× oil objective (Nikon CFI Plan Apochromat DM Lambda 100x Oil, NA = 1.45, W.D. 0.13 mm) and a ORCA-Fusion digital CMOS camera (Hamamatsu, C14440-20UP), controlled by NIS software. An automated imaging pipeline (Nikon JOBS) was used to sequentially image all wells, acquiring 8 images per well to capture at least 150 cells per condition. The images were acquired 30 min after the resuspending the cells the second time in SC media with or without glucose. During sample preparation, the timing required to image one well was accounted for to ensure that all wells were imaged at the same timepoint after treatment.

Images were analysed using Nikon NIS GA3 software and its AI modules (segment.ai, segmentobjects.ai, denoise.ai). Images were first denoised with the denoise.ai module. A subset of images from different conditions was then used to train two AI modules: one for identifying PBs and one for cell segmentation using the background mCherry signal. These modules were applied to automatically segment cells and PBs in all images and to quantify the number of PBs per cell. To remove falsely identified structures, segmented objects were filtered based on their intensity relative to the mean background intensity of their respective cells (only P bodies at least 70% brighter than the mean background were counted).

P-body counts were analysed in OriginPro 2026 (OriginLab) using the Paired Comparison Plot app. Statistical comparisons were performed using one-way ANOVA followed by Holm– Bonferroni-adjusted pairwise comparisons. The mean number of P-bodies per cell from each independent biological replicate was used as an individual observation (n = 3). Adjusted p values below 0.05 were considered statistically significant.

### Construction of yeast strains and plasmids

The strains used in this study are derivatives of W303^47,55^ and are described in Supplementary Table 6. Yeast C-terminal epitope tagging of ORFs was done by PCR-based homologous recombination^56^.

Plasmids used in this study are described in Supplementary Table 5.

For generation estradiol inducible strains, the respective plasmid carrying the lexA expression system together with the respective gene to be expressed were cut overnight by SwaI and transformed into the *dhh1Δ* strain.

### TCA protein extraction

For whole-cell protein extraction, yeast cell pellets were resuspended in 200 µL water and supplemented with 50 µL 1.875 M NaOH. Samples were vortexed and incubated on ice for 5 min. Proteins were precipitated by addition of 50 µL 100% (w/v) trichloroacetic acid (TCA). Samples were vortexed and incubated on ice for at least 10 min, followed by centrifugation at 20000 rcf for 10 min at 4 °C. The supernatant was discarded and the protein pellet was resuspended in 25 µl of 1xSDS loading buffer. 10 µL were used for Western blot analysis.

### Western blotting

Proteins were separated on NuPAGE 4–12% Bis-Tris Mini Protein Gels (1.0 mm; Thermo Fisher Scientific) at 120 V for 69 min and subsequently transferred onto PVDF membranes by semi-dry blotting in 1x Towbin buffer with 20% methanol using a Power Blotter system (Thermo Fisher Scientific). Membranes were blocked for at least 30 min in 5% (w/v) milk powder in 1x phosphate-buffered saline (PBS).

For protein detection, membranes were incubated overnight with mouse anti-V5 (GeneTex) or mouse anti-Pgk1 (Abcam) primary antibodies diluted 1:1000 in antibody dilution buffer (1x PBS, 1% [w/v] bovine serum albumin [BSA], 0.05% [w/v] sodium azide). Membranes were subsequently washed three times with approximately 25 mL washing buffer (1x PBS, 0.05% [v/v] Tween-20) and incubated for 45 min with horseradish peroxidase (HRP)-conjugated anti-mouse IgG secondary antibody (Sigma-Aldrich) diluted 1:20,000. Following secondary antibody incubation, membranes were washed extensively with washing buffer, including at least five buffer exchanges and an extended wash of at least 15 min with gentle agitation.

Chemiluminescent signals were developed using Immobilon Western chemiluminescent HRP substrate (Millipore) and detected using a Fusion FX imaging system (Vilber).

## Notes

### Competing Interest Statement

The authors have declared no competing interest.

